# Autophagic stress activates distinct compensatory secretory pathways in neurons

**DOI:** 10.1101/2024.11.07.621551

**Authors:** Sierra D. Palumbos, Jacob Popolow, Juliet Goldsmith, Erika L.F. Holzbaur

**Affiliations:** Department of Physiology, Perelman School of Medicine, University of Pennsylvania, Philadelphia, PA 19104, USA; Aligning Science Across Parkinson’s (ASAP) Collaborative Research Network, Chevy Chase, MD 20815, USA

**Keywords:** Autophagy, Parkinson’s disease, Secretion, Mitochondria, Neurodegeneration

## Abstract

Autophagic dysfunction is a hallmark of neurodegenerative disease, leaving neurons vulnerable to the accumulation of damaged organelles and proteins. However, the late onset of diseases suggests that compensatory quality control mechanisms may be engaged to delay the deleterious effects induced by compromised autophagy. Neurons expressing common familial Parkinson’s disease (PD)-associated mutations in LRRK2 kinase exhibit defective autophagy. Here, we demonstrate that both primary murine neurons and human iPSC-derived neurons harboring pathogenic LRRK2 upregulate the secretion of extracellular vesicles. We used unbiased proteomics to characterize the secretome of LRRK2^G2019S^ neurons and found that autophagic cargos including mitochondrial proteins were enriched. Based on these observations, we hypothesized that autophagosomes are rerouted toward secretion when cell-autonomous degradation is compromised, likely to mediate clearance of undegraded cellular waste. Immunoblotting confirmed the release of autophagic cargos and immunocytochemistry demonstrated that secretory autophagy was upregulated in LRRK2^G2019S^ neurons. We also found that LRRK2^G2019S^ neurons upregulate the release of exosomes containing miRNAs. Live-cell imaging confirmed that this upregulation of exosomal release was dependent on hyperactive LRRK2 activity, while pharmacological experiments indicate that this release staves off apoptosis. Finally, we show that markers of both vesicle populations are upregulated in plasma from mice expressing pathogenic LRRK2. In sum, we find that neurons expressing pathogenic LRRK2 upregulate the compensatory release of secreted autophagosomes and exosomes, to mediate waste disposal and transcellular communication, respectively. We propose that this increased secretion contributes to the maintenance of cellular homeostasis, delaying neurodegenerative disease progression over the short term while potentially contributing to increased neuroinflammation over the longer term.

**SIGNIFICANCE:** A hallmark feature of many neurodegenerative diseases is autophagy dysfunction, resulting in the accumulation of damaged proteins and organelles that is detrimental to neuronal health. The late onset of neurodegenerative diseases, however, suggests alternative quality control mechanisms may delay neuronal degeneration. Here, we demonstrate that neurons expressing a Parkinson’s Disease-causing mutation upregulate the release of two extracellular vesicle populations. First, we observe the increased expulsion of secreted autophagosomes to mediate cellular waste disposal. Second, we observe the increased release of exosomes, likely to facilitate transcellular communication. Thus, we propose that increases in secretory autophagy and exosome release are a homeostatic response in neurons undergoing chronic stress.

## INTRODUCTION

Macroautophagy, hereafter referred to as autophagy, is a fundamental cellular process by which aggregated proteins and dysfunctional organelles are degraded (1). Neurons utilize autophagy as a homeostatic mechanism, exhibiting robust basal autophagy in the absence of cellular stressors. Knockout of critical components of the autophagy machinery, including Atg5 and Atg7 is sufficient to induce neurodegeneration (2, 3). In neurons under basal conditions, cargos to be cleared by autophagy are captured within double membrane autophagosomes that forms preferentially at axonal termini as well as presynaptic sites and are then trafficked back toward the cell soma (4, 5). Autophagosomes fuse with lysosomes en route to the soma in order to degrade internalized cargos (6). Disruption of either autophagosome trafficking or lysosomal fusion inhibit compartment maturation and the subsequent degradation of cargos internalized within axonal autophagosomes (6, 7).

In many neurodegenerative diseases, autophagic dysfunction results in the accumulation of protein aggregates and dysfunctional organelles, threatening neuronal homeostasis (1, 8, 9). Parkinson’s Disease (PD), for example, is a progressive neurodegenerative disease characterized by autophagic dysfunction, accumulation of α-synuclein aggregates, and neuroinflammation (9–11). Mutations that induce hyperactivity of the kinase LRRK2, including LRRK2^G2019S^ and LRRK2^R1441H^, represent some of the most common causes of familial PD. LRRK2 hyperactivity leads to aberrant phosphorylation of Rab GTPases, key regulators of trafficking pathways in the cell (12, 13), including the trafficking and maturation of neuronal autophagosomes (**Figure 1A**) (7, 14–16).

**Figure 1:**
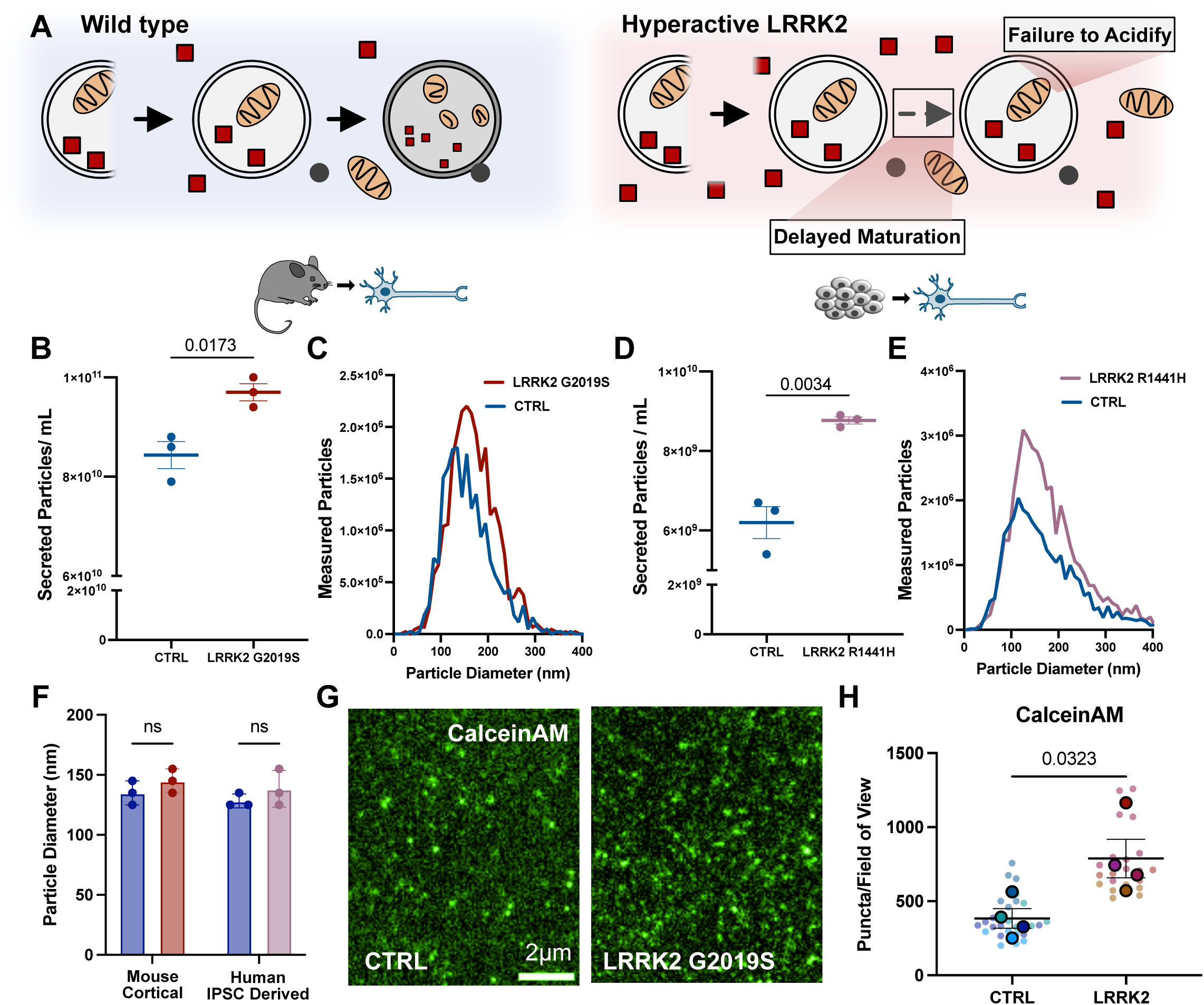
Neurons expressing hyperactive LRRK2 upregulate secretion. **A)** Schematic representing degradative-autophagy in control (left) and LRRK2 mutant (right) neurons. Neurons expressing hyperactive LRRK2 exhibit delayed autophagosome trafficking down the axon, stalled autophagosome maturation and reduced acidification. **B)** Quantification of Nanoparticle Tracking Analysis (NTA) of secreted particles isolated from DIV11 wild type and LRRK2^G2019S^ primary cortical neurons. Particle count normalized to cell number. N=3, two-tailed t-test, error bars represent mean and SEM. **C)** Representative size distribution of measured particles released from wild type and LRRK2^G2019S^ primary cortical neurons in individual experiment. **D)** Quantification of NTA of secreted particles isolated from DIV21 wild type and LRRK2^R1441H^ human iPSC derived KOLF2.1J neurons. Particle count normalized to cell number. N=3, two-tailed t-test, error bars represent mean and SEM. **E)** Representative size distribution of measured particles released from control and LRRK2^R1441H^ human iPSC derived KOLF2.1J neurons in an individual experiment. **F)** Measured particle diameter of the most common particle size (mode) from control and LRRK2 neurons in both mouse primary cortical neurons and human iPSC derived KOLF2.1J neurons. N=3, 2-way ANOVA with uncorrected Fisher’s least significant difference (LSD), error bars represent mean and SEM. **G)** Representative TIRF images of vesicles treated with Calcein AM isolated from wild type (left) and LRRK2^G2019S^ (right) primary cortical neurons. Images have been pseudo-colored. **H)** Superplot representing measured CalceinAM fluorescing puncta per field of view of vesicles isolated from control and LRRK2^G2019S^ primary cortical neurons. N=4, two-tailed t-test comparing biological replicates, error bars represent SEM.

We hypothesized that the resulting strain on autophagy-dependent degradation in neurons might induce activation of alternative quality control mechanisms to prevent or delay proteotoxicity. One possibility is that neurons expressing hyperactive LRRK2 may upregulate secretion, ejecting undegraded cellular waste to the extracellular space via autophagy-dependent secretion. Secretory autophagy is a broad term encompassing multiple pathways that rely on autophagic proteins to mediate the release of diverse contents (17, 18). In canonical secretory autophagy, autophagic proteins mediate the fusion of the outer membrane of an autophagosome with the plasma membrane to release soluble proteins trapped between the vesicular membranes and the inner autophagic vesicle (19, 20). Non-canonical secretory autophagy pathways have also been identified that mediate the release of different vesicle populations or organelles, including exosomes (21), exophers (22), and mitochondria (23). Given that inhibition of autophagosome-lysosome fusion has been shown to trigger the upregulation of autophagy-dependent secretion (24–27), we reasoned that neurons expressing pathogenic LRRK2 mutations may similarly engage secretion as a compensatory mechanism in response to autophagic dysfunction.

Here, we demonstrate that neurons expressing hyperactive LRRK2 activate the compensatory release of both secreted autophagosomes and exosomes. We performed unbiased proteomic and transcriptomic characterization of vesicles released from LRRK2^G2019S^ neurons, demonstrating that secretory autophagy exports known cargos of autophagy, including mitochondria and synaptic proteins, while we find that the secreted exosomes contain microRNAs (miRNAs) known to regulate autophagy and inflammation. As further confirmation of our model, we used both fixed and live-cell imaging to demonstrate that the two vesicle populations exhibit distinct spatial and temporal dynamics of release. Further, we find that the release of exosomes is neuroprotective. Finally, we show that release of both vesicle populations is upregulated in plasma of a mouse line expressing pathogenic LRRK2^G2019S^. Based on these findings, we propose that activation of these two secretion pathways mediate distinct roles, the release of cellular waste via secreted autophagosomes and transcellular communication via exosomes. Together, we propose that upregulation of these secretory pathways acts as a compensatory mechanism to sustain cellular homeostasis when autophagy-mediated degradation is impaired. Importantly, however, the elevated secretion of proinflammatory contents such as mitochondrial DNA (28, 29) and miRNAs (30, 31) is likely to negatively affect neuronal health over the longer term, contributing to neurodegenerative disease progression in patients with PD.

## RESULTS

### LRRK2 mutant neurons upregulate secretion

Pathogenic mutations in LRRK2 induce kinase hyperactivity, leading to inappropriate phosphorylation of kinase substrates, predominantly organelle-associated Rab GTPases (15), and a resulting disruption in intracellular trafficking pathways (12), specifically disrupting autophagosome maturation in neurons (7, 16). To test our hypothesis that this disruption might lead to a compensatory increase in secretion, we began by focusing on the most common PD-linked mutation in LRRK2, a G2019S missense mutation. We isolated embryonic cortical neurons from wild type and LRRK2^G2019S^ mice. Conditioned media were collected from cultured neurons over 10 days and EVs were enriched from the harvested media. For each replicate, the same number of neurons were plated, and the resulting EV measurements were normalized total protein levels in cell lysates. We tracked the size and concentration of particles secreted using Nanoparticle Tracking Analysis (NTA) and observed a 15% increase in particles in EV samples isolated from LRRK2^G2019S^ neurons as compared to EV samples isolated from control neurons (**Figure 1B, C)**. No significant changes in size distribution were detected, with the most commonly detected particle size being 135nm in wild type and 145nm in LRRK2^G2019S^ samples (**Figure 1F**). Several shoulder peaks were also detected in our analysis of both wild type and LRRK2 particles, suggesting the release of multiple vesicle populations over a range of sizes (**Figure 1C)**.

To extend this observation, we examined EV secretion from human iPSC-derived neurons (KOLF2.1J (32)) harboring a distinct Parkinson’s Disease-causing mutation, LRRK2^R1441H^. This mutation results in a more profound hyperactivation of the LRRK2 kinase as compared to G2019S, leading to more profound deficits in autophagosome trafficking (15, 16). Again, we observed significantly more particles released from LRRK2^R1441H^ neurons as compared to isogenic control neurons (a 42% increase) (**Figure 1D),** with no change in size distribution detected between the genotypes **(Figure 1F**). While this is a cross-species comparison, the effect size on secretion appears to be correlated with the extent of kinase hyperactivation.

To confirm these findings, we used CalceinAM to assay intact vesicles collected from WT and LRRK2^G2019S^ neurons following EV isolation via ultracentrifugation (33). CalceinAM fluoresces after passively entering an EV that is unpermeabilized and thus the fluorescent signal corresponds to intact, membrane-enclosed structures. Consistent with our NTA results, we observed significantly more CalceinAM puncta in samples collected from LRRK2^G2019S^ mutant neurons (**Figure 1G, H),** indicating increased secretion of extracellular vesicles. Thus, in both mouse and human neurons, hyperactive LRRK2 induces increased secretion of extracellular vesicles.

### Proteomic profiling of offloaded vesicles

Extracellular vesicles serve a variety of purposes (34, 35), including waste disposal (22, 36) and transcellular signaling (37), and therefore may contain a wide range of internalized cargos and transmembrane proteins that can vary by vesicle class and cell type. To characterize the released vesicles, we first performed quantitative proteomic analysis of isolated vesicles from 5 independent cultures of control and of LRRK2^G2019S^ primary cortical neurons. Extracellular vesicles were enriched via ultracentrifugation to broadly capture released EVs. The protein concentration of isolated EVs was normalized across samples prior to quantitative mass spectrometry analysis (**Supplemental Figure 1A-B**). Detected proteins were assigned an expression value based on the number of detected reads within a given sample (median expression value = 0). Vesicle enrichment for both genotypes was confirmed by comparison to previous EV proteomic analysis that characterized proteins found in either small or large vesicle fractions (38) (**Figure 2A, Supplemental Figure 1C**). We noted the presence of 89/100 of the most common EV-associated proteins, which were readily detected above the median detection threshold in both control and LRRK2 samples (expression value > 0) (39) (**Figure 2E, Supplemental Figure 1D**). Next, we compared the expression of detected proteins associated with either small EVs (**Figure 2B**) or large EVs (**Figure 2C**) across genotypes. While we detected no change in the average expression value of small-EV associated proteins, we did detect significantly higher expression values for large-EV associated proteins in LRRK2^G2019S^ samples, suggesting that large EVs were likely enriched in these samples.

**Figure 2:**
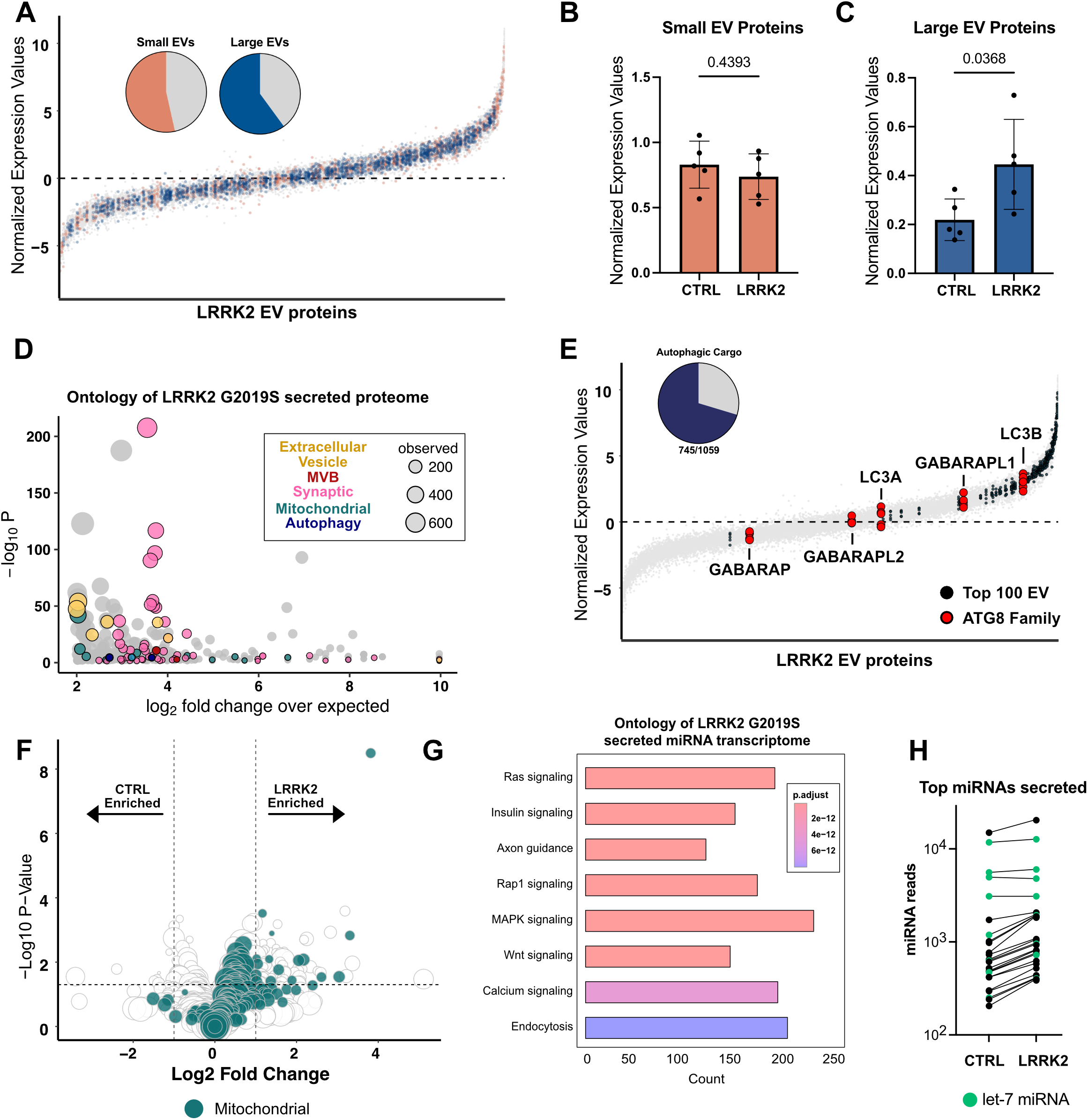
Proteomic profiling reveals autophagic cargo secreted from LRRK2^G2019S^ primary cortical neurons. **A)** Percent of proteins (pie chart inserts) characterized as “Small EV enriched” or “Large EV enriched” by Lischnig et al.(38), detected in LRRK2 secretome. Proteins detected in extracellular vesicles isolated from 5 replicate samples of LRRK2^G2019S^ neurons ranked by abundance. Median protein abundance = 0. Large EV proteins colored blue, Small EV proteins colored orange. **B)** Average expression value of all detected Small EV(38)proteins between control and LRRK2 ^G2019S^ secretome across replicates. N=5, two-tailed t-test, error bars represent SEM. **C)** Average expression values of all detected Large EV(38)proteins between control and LRRK2 ^G2019S^ secretome across replicates. N=5, two-tailed t-test, error bars represent SEM. **D)** Bubble plot representation of ontology terms of top 50% of detected proteins in LRRK2^G2019S^ secretome. Ontology analysis of GO cellular component by PANTHER. Each bubble depicts unique GO term and size of bubble represents number of proteins within term that was detected. Bubbles are pseudo coated by terms indicated in box inset. **E)** Proteins detected in LRRK2^G2019S^ secretome ranked by abundance. Black dots indicate proteins defined as top 100 EV-associated cargo. Red dots indicate members of ATG 8 family. Percent of proteins characterized as autophagic cargo(40) (pie chart insert) detected in LRRK2^G2019S^ EVs. **F)** Volcano plot representing differentially secreted proteins from control and LRRK2^G2019S^ neurons. Green circles indicate mitochondrial associated proteins as defined by MitoCarta 3.0. Size of dot correlates to normalized expression value. **G)** Bubble plot representation of ontology terms from proteins differentially expressed between control and LRRK2^G2019S^ EVs. Significance threshold determined by p-value of differential abundance analysis. Each bubble depicts unique GO term. Bubbles are pseudo colored based on broader terms indicated below. Size of bubble represents number of proteins representing that term.

Ontology analysis of the secretomes of LRRK2 ^G2019S^ and wild type confirmed that proteins associated with secretion, including those associated with exosome formation, e.g., multivesicular body (MVB) proteins, were readily released (**Figure 2D and Supplemental Figure 1E**). Interestingly, we noted that proteins associated with synaptic function, mitochondria and autophagy were also enriched in the secretomes from both LRRK2 ^G2019S^ and wild type neurons (**Figure 2D and Supplemental Figure 1E**). Recently, we used proteomic analysis to define the basal cargos of neuronal autophagosomes and detected both synaptic proteins and mitochondria as enriched autophagosome cargo (40). Given this commonality, we reasoned that these neurons could be releasing a population of autophagic vesicles via secretory autophagy. To test this possibility, we compared all detected proteins in EVs isolated from LRRK2^G2019S^ or control neurons to the previously characterized proteome of autophagosomal cargos from brain (40). In both cases, over 70% of known autophagic cargos from brain were detected within our EV proteomic datasets (**Figure 2E**).

If wild type and LRRK2^G2019S^ neurons are releasing secreted autophagosomes, members of the ATG8 family, well-established markers of autophagosomes, should be detected among the secreted proteins. Consistent with this, nearly all secretory autophagy pathways have reported the release of the autophagic protein LC3B (17, 21, 22, 24, 25), a member of the ATG8 family. In line with previous work, we detected 5 of 6 ATG8s in both the LRRK2^G2019S^ and control samples, with the highest detected being LC3B (**Figure 2E, Supplemental Figure 1D**). Thus, unbiased proteomic profiling of the secretomes from either control or LRRK2^G2019S^ primary cortical neurons detected markers associated with multiple classes of extracellular vesicles, including exosomes and secreted autophagosomes.

To determine how the secretory proteome was altered in LRRK2^G2019S^ mutants, we performed differential expression analysis and observed 250 upregulated proteins and 80 downregulated proteins in samples from LRRK2^G2019S^ neurons as compared to control neurons (**Figure 2F**). Ontology analysis of the differentially expressed proteins suggests that mitochondrial proteins are the most prominent cargo upregulated in LRRK2^G2019S^ EVs (**Supplemental Figure 1F**). Indeed, 92/250 upregulated secreted proteins are classified as mitochondrial by MitoCarta3.0(41). This observation could indicate that secreted autophagosomes are more readily released by LRRK2^G2019S^ as 20% of all neuronal autophagosome cargo is mitochondrially-derived (40). The contents of LRRK2^G2019S^ autophagosomes are largely unchanged from wild type, with mitochondria being readily targeted for autophagosome engulfment (42). Further, while autophagosome maturation is impaired in LRRK2^G2019S^ neurons, overall numbers of autophagosomes are not significantly impacted. Thus, our findings suggest that LRRK2^G2019S^ neurons, which exhibit autophagosome maturation defects, may be redirecting unacidified autophagosomes toward extracellular release.

### Transcriptomic profiling of released vesicles

In addition to proteins, many extracellular vesicles contain RNAs, including miRNAs, to mediate transcellular communication. We therefore performed unbiased small RNA (smRNA) sequencing of isolated vesicles from both LRRK2^G2019S^ and wild type neurons. Secreted extracellular vesicles were isolated via ultracentrifugation followed by RNA extraction, smRNA library preparation, and subsequent sequencing of the normalized libraries. In EVs isolated from both wild type and LRRK2^G2019S^, miRNAs were detected, with 391 and 499 miRNAs reaching our detection threshold (average of at least 2 reads across 5 samples), respectively. KEGG analysis of the downstream targets of the top 25 miRNAs detected in EVs isolated from both wild type and LRRK2^G2019S^ neurons confirmed that the secreted vesicles were likely mediating transcellular communication (**Figure 2G**). Many signaling pathways, including ones that impact inflammatory responses, e.g., MAPK, were enriched in both the wild type and LRRK2^G2019S^ secretome (43). In both genotypes, 4 of the top 5 detected miRNAs were of the *let-7* miRNA family (**Figure 2H**), which play a role in many processes, including autophagy and inflammation (44, 45). Thus, both wild type and LRRK2^G2019S^ neurons are offloading miRNAs which could be relevant to neuronal homeostasis. We next asked whether any miRNAs were differentially expressed, and detected no miRNAs significantly upregulated in either genotype (**Supplemental Figure 2A**). While no individual miRNA was significantly upregulated in LRRK2 EVs, the average levels of the top 50 detected miRNAs were all elevated in LRRK2 compared to wild type (**Figure 2I, Supplemental Figure 2B**). Together, our immunoblots and transcriptomics results argue that it is the overall quantity, and not the contents, of released exosomes that are altered in LRRK2^G2019S^ neurons.

### Hyperactive LRRK2 enhances the secretion of distinct classes of extracellular vesicles

To identify the class(es) of extracellular vesicles being offloaded from LRRK2^G2019S^ neurons, we performed differential ultracentrifugation of conditioned media to separate EV populations based on density. Large extracellular vesicles, including microvesicles and secreted autophagosomes, were enriched via a 20,000xg spin (large EVs/P20), followed by sequential enrichment of small EVs, including exosomes, via a 100,000xg spin (small EVs/P100) (**Figure 3A**)(46). Large (P20) and small (P100) EV pellets were isolated and analyzed by immunoblot using antibodies against known EV-associated transmembrane proteins and cargos (**Figure 3B**). Given that our proteomics results closely mirrored previous proteomic profiling of brain autophagosomes, we first probed for LC3/ATG8 which is offloaded in many secretory autophagy pathways. LC3-I is conjugated to phosphatidylethanolamine to form LC3-II, which marks the forming autophagosome and stays associated throughout autophagosome maturation (47, 48). Across both genotypes, the majority of both LC3-I and LC3-II was detected in the large EV/P20 fraction. Of note, we observed significantly more LC3-II in EVs isolated from LRRK2^G2019S^ neurons as compared to control neurons in the P20 fraction (2.1-fold increase, p=0.0089) (**Figure 3C, D**).

**Figure 3:**
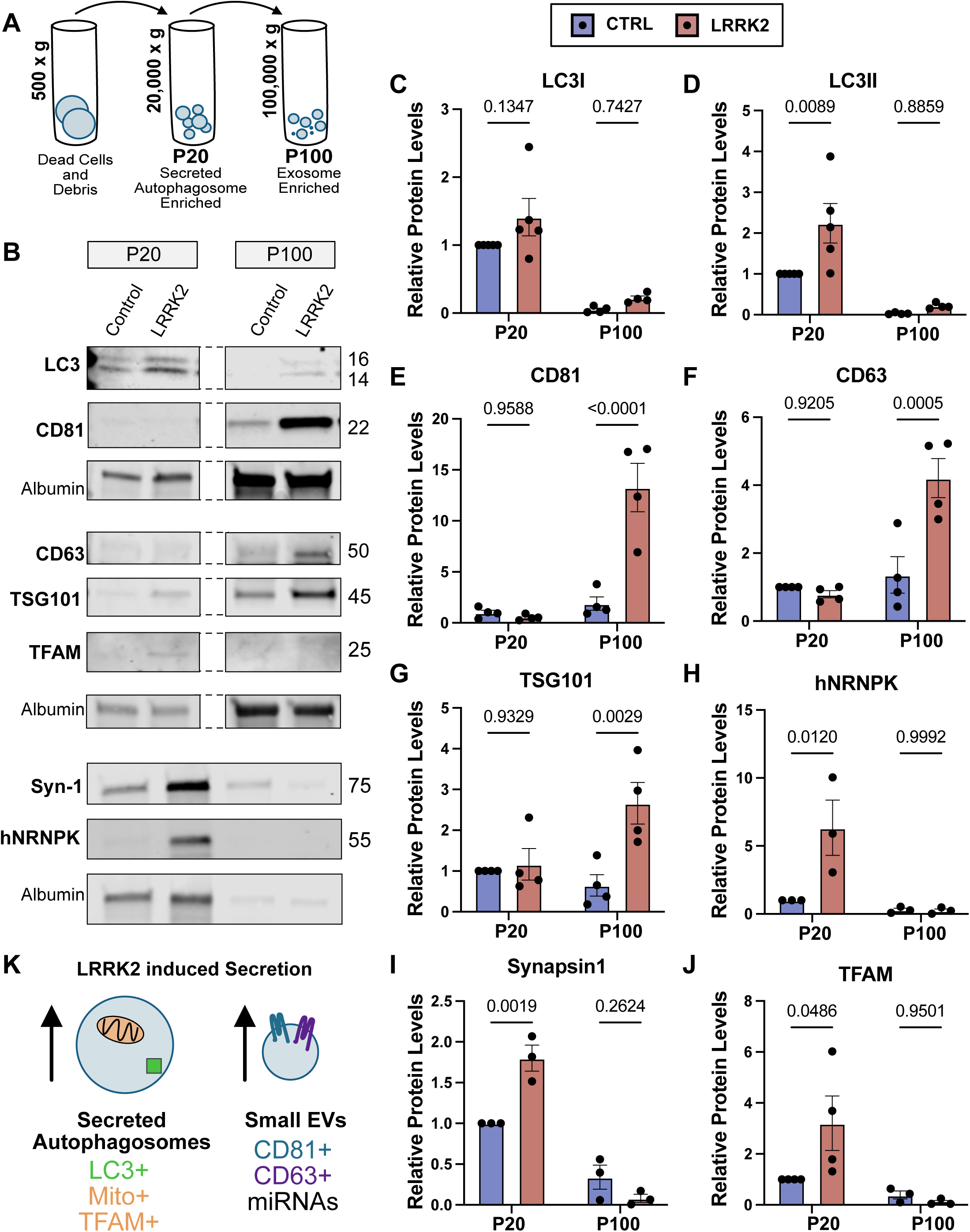
Two secretory pathways are upregulated in LRRK2^G2019S^ neurons. **A)** Schematic representing differential ultracentrifugation approach to enrich for secreted autophagosomes and exosomes. P20 indicates pellet isolated following 20,000g spin and P200 indicates pellet isolated following 100,000g spin. Additional vesicle populations are captured with this approach. **B)** Representative western blots of isolated P20 and P100 fractions from primary cortical control and LRRK2^G2019S^ neurons. Dashed lines indicate corresponding lanes are from the same gel. Detected protein and corresponding molecular weight indicated. Equal volume was loaded for each well and samples were normalized to amount of detected media protein, albumin. **C-J)** Quantifications of relative levels of detected band intensities for **C)** LC3-I, **D)** LC3-II, **E)** CD81, **F)** CD63, **G)** TSG101, **H)** hNRNPK, **I)** Synapsin-1, and **J)** TFAM from P20 and P100 vesicular fractions isolated from control (blue) and LRRK2^G2019S^ (red) neurons. All quantifications of band intensities are normalized to the control P20 band to clearly indicate relative abundance of detected proteins. Individual replicates represented by black dots. 2way ANOVA with Šídák’s multiple comparison test. Error bars indicate SEM. **K)** KEGG analysis of targets of the top miRNAs detected in both wild type and LRRK2^G2019S^ neuronal secretome. The top 25 miRNAs was consistent between both genotypes. **L)** Average number of reads of top 25 miRNAs detected in control and LRRK2^G2019S^ secreted transcriptome. For each miRNA, read average is elevated in LRRK2^G2019S^. In both genotypes, 4 of the top 5 miRNAs are members of the let-7 family. **M)** Schematic representing two vesicle populations that are upregulated in LRRK2^G2019S^ neurons. Presumed secreted autophagosomes (left) contain LC3-II, mitochondria, synaptic proteins and RNA-binding proteins. Presumed exosomes express CD81, CD63 and TSG101.

Most extracellular vesicles are enriched for a family of transmembrane proteins, the tetraspanins, which form microdomains on vesicles important for functions including biogenesis and cargo sorting. We compared the expression of CD81 and CD63, the two most highly enriched tetraspanins on exosomes (49), on EVs isolated from control vs. LRRK2^G2019S^ neurons and observed no change in CD81 or CD63 expression in the large EV/P20 fraction. Thus, the large LC3-II+ vesicles being secreted from LRRK2 mutant neurons lack tetraspanins, an observation consistent with their identification as secreted autophagosomes. Strikingly, in the P100 fraction enriched for small EVs, we observed significantly more CD81 (8.4 fold increase) and CD63 (2.6 fold increase) in EVs released from LRRK2 mutant neurons as compared to wild type neurons (**Figure 3E, F**). Changes in the relative enrichment of these proteins were not due to changes in protein expression as no significant changes were observed by immunoblot analysis of neuronal lysates (**Supplemental Figure 3**).

Together, these observations suggest that two populations of EVs are upregulated in LRRK2 mutants: 1) larger EVs that are positive for LC3-II but lack canonical tetraspanins and 2) small EVs enriched for CD63 and CD81. Given our proteomic observations suggesting that autophagic cargo are expelled from LRRK2 neurons and previous literature that has found CD63 expression largely restricted to exosomes, we propose that these two EV populations (large EVs and small EVs) represent secreted autophagosomes and exosomes, respectively.

### Large and small EVs contain distinct cargos

To provide additional insight into the nature and function of the two populations of secreted vesicles detected, we used immunoblotting to compare cargos associated with either secreted autophagosomes or exosomes. Given the size and density of autophagosomes (50), we first asked if known autophagosome cargos were enriched in our P20 fraction. Previously, our lab defined the contents of neuronal autophagosomes under basal conditions and found that both synaptic proteins, including Synapsin-1, and mitochondria enriched in the nucleoid marker TFAM are targeted to autophagosomes under basal conditions (40). More recently, we found that several cargos are increased in autophagosomes isolated from the brains of mice expressing mutant LRRK2, including the RNA-binding protein, hnRNPK (42). hnRNPK has been shown to interact with LC3 and is secreted within EVs released via LC-3 Dependent Extracellular vesicle Loading and Secretion (LDELS)(25). Consistent with our model that hyperactive LRRK2 activity induces the release of secreted autophagosomes, we saw significant increases in the release of Syn-1 (1.8-fold increase), TFAM (3.2-fold increase), and hnRNPK (5.9-fold increase) in the LRRK2 P20 pellet as compared to controls when measured by immunoblotting (**Figure 3 G-I**). In contrast, these proteins were minimally detected in the P100 fractions from either LRRK2^G2019S^ or wild type neurons.

Exosomes originate in the multivesicular body, or MVB, where different contents, e.g., ESCRT proteins and RNAs, are loaded into small vesicles before being released into the extracellular space (51). We tracked the expression of the ESCRT protein and canonical EV cargo, TSG-101 and observed a 3.5-fold increase in the small EV/P100 fraction in LRRK2^G2019S^ mutants (**Figure 3H**). In contrast, we observed no significant change in enrichment of TSG-101 in the P20 fraction corresponding to large EVs. Exosomes are very small and thus contain limited cargo, often enriched for miRNA and mRNA (51, 52). Our smRNA transcriptomic analysis confirmed that miRNAs were released by both wild type and LRRK2^G2019S^ neurons. To determine if RNAs were contained within secreted exosomes, we asked whether P100 fractions isolated from wild type or LRRK2^G2019S^ neurons contained RNA. We used the RNA specific dye, SYTO RNASelect (Thermo), to measure relative levels in fluorescence across our different conditions and observed that RNA could be readily detected in both wild type and LRRK2^G2019S^ P100 fractions (**Supplemental Figure 2C**), suggesting that RNAs are likely cargos within released exosomes.

### LRRK2 mutant neurons release secreted autophagosomes

ATG8 family members (LC3/GABARAP) are stably associated with both the inner and outer membrane of mature autophagosomes. In canonical secretory autophagy, the outer LC3/GABARAP membrane fuses with the plasma membrane to release both lumen contents and the inner autophagosome vesicle (53). If autophagosome content is secreted in this manner from the cortical neurons examined here, LC3 should be detected at the cell surface. We transfected wildtype and LRRK2^G2019S^ neurons with fluorescently labeled LC3 and tracked expression at the cell surface using live TIRF microscopy. Consistent with secretory autophagy events, we observed frequent LC3 bursts at the cell surface in both control and LRRK2^G2019S^ neurons (**Figure 4A-B**). To quantitate this effect, we measured external LC3/GABARAP by immunocytochemistry using an antibody against GABARAPL1/L2/L3 in unpermeabilized control (**Figure 4C**) and LRRK2^G2019S^ neurons (**Figure 4E**). We detected significantly more GABARAPL1/L2/L3 at the cell surface of LRRK2^G2019S^ neurons, consistent with a higher level of secretory autophagy (**Figure 4G**). We confirmed that this signal could be detected in TIRF (**Figure 4D, F**) and was distinct from LC3/GABARAP localization in permeabilized cells (**Supplemental Figure 3B**).

**Figure 4:**
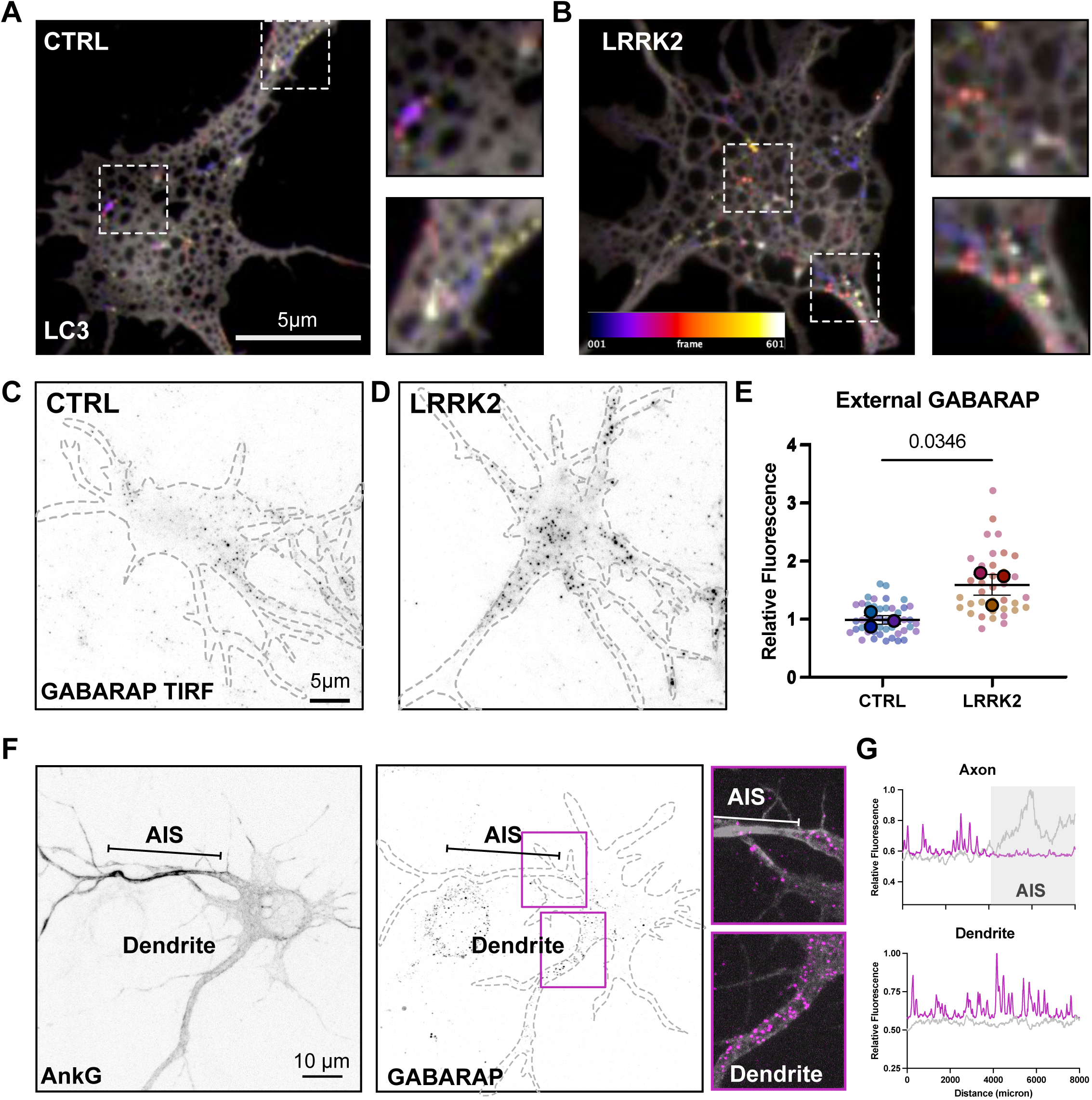
Secreted autophagosomes are upregulated in LRRK2^G2019S^ neurons. **A-B)** Representative frame sequence of **A)** control and **B)** LRRK2^G2019S^ neurons transfected with LC3 mScarlet captured with TIRF microscopy over 5-minute period. Temporal color code represents frames from 1-601 (2 frames/sec). Increases in fluorescence intensity at cell surface are consistent with secretory autophagy events. Dashed boxes indicate example LC3 fluorescent event on soma and individual process. **C-D)** Representative TIRF imagee of external GABARAP signal in **C)** wild type and **D)** LRRK2^G2019S^ neurons. **E)** Quantification of external GABARAP signal. Superplot of biological and technical replicates. N=3, two-tailed t-test comparing biological replicates, error bars represent SEM. **F)** Representative image LRRK2^G2019S^ neuron transfected with AnkyrinG and immunstained for external GABARAP. Axon Initiation Segment (AIS) indicated by bar. Insets of AIS and dendrite. **G)** Representative linescans of GABARAP (magenta) and AnkG (grey) of axon and dendrite of images from F.

Interestingly, we noted that in both our live and fixed imaging approaches, surface GABARAPL1/L2/L3was spatially enriched in a somal compartment which often extended into a single tapered process, consistent with the primary dendrite (**Figure 4C-E**). Morphological characterization confirmed that in ∼70% of neurons imaged, an accumulation of external GABARAPL1/L2/L3 could be observed at the presumed primary dendrite. We transfected neurons with the AIS marker, Ankyrin G (AnkG) and confirmed that the accumulation of GABARAPL1/L2/L3 was dendritic, rather than axonal (**Figure 4F)**. Thus, LC3 secretion events are restricted to the somatodendritic compartment. This observation mirrors previous observations from live cell imaging indicating that once axonal autophagosomes enter the soma, they are constrained to the somal and dendritic regions(5).

### LRRK2 mutant neurons upregulate release of exosomes

Our immunoblotting results suggest that small EVs containing both CD81 and CD63 are also secreted from LRRK2^G2019S^ neurons. As immunoblotting relies on the pooling of EVs to look for global changes, we next used ExoView (Spectradyne) to track tetraspanin expression at the level of individual EVs. We captured EVs from both wild type and LRRK2^G2019S^ neurons using an anti-CD81 antibody followed by subsequent antibody detection against CD81, CD9 and CD63 (**Figure 5A**). Consistent with our earlier observations, more CD81+ EVs were released from LRRK2^G2019S^ neurons compared to wild type neurons (**Figure 5B**). Additionally, we detected more CD81+/CD63+, and CD81+/CD9+ dual-labeled vesicles captured from LRRK2^G2019S^ EVs (**Figure 5B**). These observations corroborated our immunoblotting results, where we observed substantial CD81 and CD63 release from LRRK2^G2019S^ neurons, consistent with the release of exosomes.

**Figure 5:**
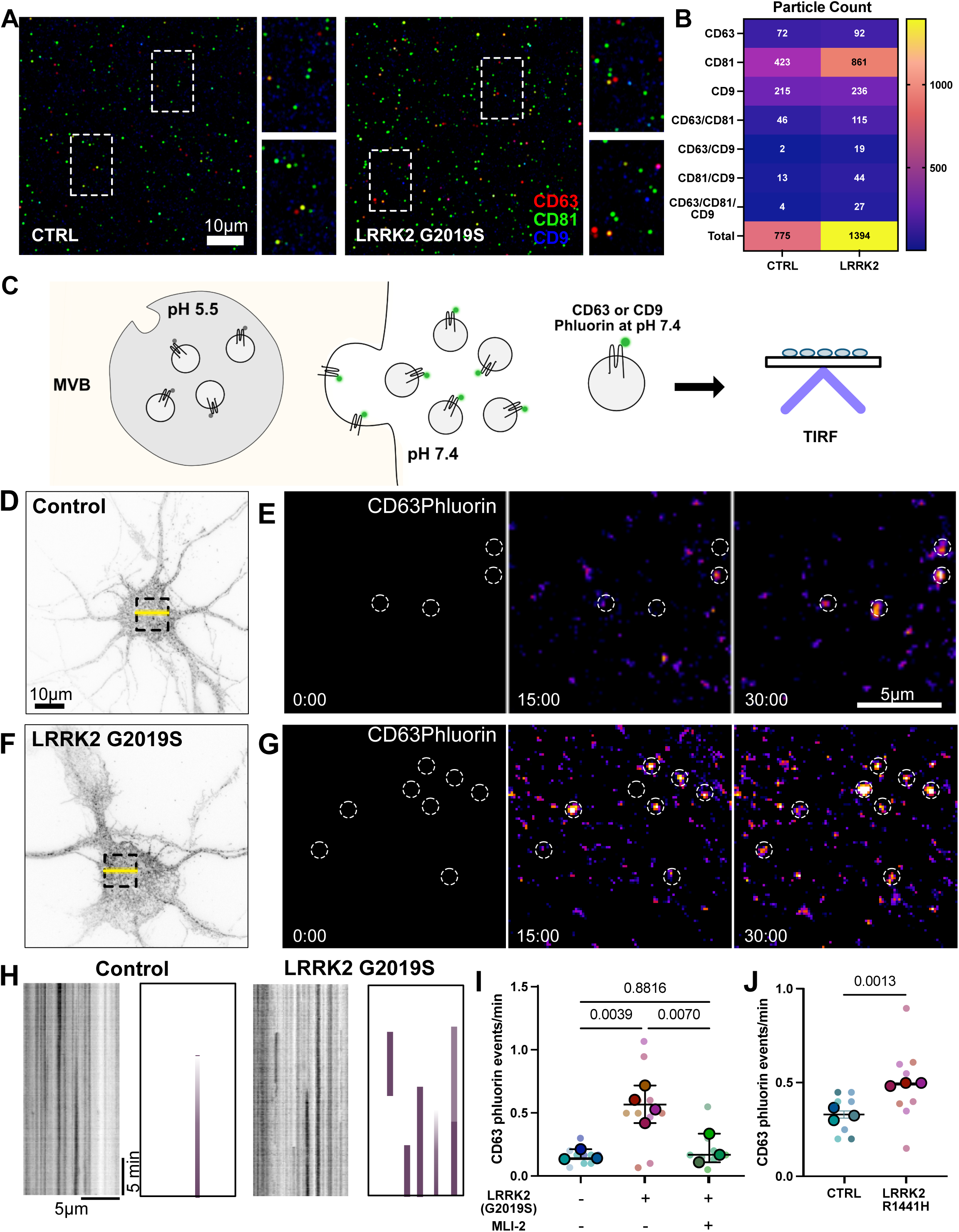
Secretion of exosomes is augmented in LRRK2^G2019S^ neurons. **A)** Representative image of ExoView anti-CD81 captured vesicles from control and LRRK2^G2019S^ neurons. Zoomed in views indicated by dashed rectangles. **B)** Quantification of detected particles positive for individual or combination of tetraspanins, CD81, CD63, and CD9. Total particle count noted. **C)** Schematic indicating imaging pipeline to detect CD63pHluorin secretion events. CD63pHluorin signal is quenched in acidic environments. When the multivesicular body (MVB) fuses with the plasma membrane, CD63pHluorin will fluoresce. Neurons expressing CD63pHluorin were imaged in TIRF microscopy to capture secretion events at the cell surface. **D)** Representative image of CD63pHluorin expressing control neuron. Dashed box indicates panels depicted in panel E. Yellow line indicates kymograph depicted in panel H (left). **E)** Time series of CD63pHluorin expression in control neuron. The first frame was subtracted from all subsequent frames. Time stamp indicated in each frame. Dashed circles indicate fusion events. **F)** Representative image of CD63pHluorin expressing LRRK2^G2019S^ neuron. Dashed box indicates panels depicted in panel G. Yellow line indicates kymograph depicted in panel H (right). **G)** Time series of CD63pHluorin expression in LRRK2^G2019S^ neuron. The first frame was subtracted from all subsequent frames. Time stamp indicated in each frame. Dashed circles indicate fusion events. **H)** Representative kymographs and schematic indicating CD63pHluorin fusion events over time. **I)** Quantification of CD63pHluorin events in control, LRRK2^G2019S^, and LRRK2^G2019S^ + MLI-2 primary cortical neurons. Superplot indicating biological and technical replicates. N=4, ordinary one-way ANOVA with Tukey’s multiple comparison test of biological replicates, error bars represent SEM. **J)** Quantification of CD63pHluorin events in control, LRRK2^R1441H^ KOLF2.1J neurons. Superplot indicating biological and technical replicates. N=3, two-tailed t-test comparing biological replicates, error bars represent SEM.

CD63 is a commonly used marker of exosomes (51, 54). Given our immunoblot results that CD63 is significantly enriched in the small EV/P100 fraction from LRRK2^G2019S^ neurons, we sought to better characterize the secretion of CD63 positive EVs using live cell imaging. We expressed the pH sensitive fluorophore, pHluorin, tagged to CD63 (CD63-pHluorin) in primary cortical neurons isolated from wild type or LRRK2^G2019S^ mice. The modified GFP signal of CD63pHluorin is quenched in acidic environments such as the MVB, and is only observed upon fusion with the extracellular membrane for EV release (**Figure 5C**). We measured CD63-pHluorin signal in control and LRRK2^G2019S^ neurons using TIRF microscopy. New fusion events could be readily identified when the first frame of the video was subtracted from all subsequent frames (**Figure 5D-G**), or via kymographs (**Figure 5H**). We observed a 4-fold increase in fusion events in LRRK2^G2019S^ neurons compared to wild type neurons suggesting that exosomal secretion is upregulated in LRRK2^G2019S^ neurons (**Figure 5I**). Additionally, this upregulation of CD63 secretion events is dependent on LRRK2 hyperactivity as pre-treatment of neurons with MLI-2, a selective LRRK2 kinase inhibitor(55), blocked the observed increase of CD63 fusion events (**Figure 5I**). In contrast to the LC3 secretion events described above, CD63-postitive release events were primarily observed on the cell soma with no clear spatial clustering near or at the primary dendrite, although occasionally fusion events could be observed on neurites. Individual fusion events could be observed to persist beyond the length of the movie, suggesting that CD63 could be docked upon MVB fusion as well as released.

To determine whether human neurons expressing the LRRK2^R1441H^ mutation similarly released CD63+ EVs, we also performed live cell imaging of LRRK2^R1441H^ and isogenic control iPSC-derived neurons expressing CD63-pHluorin. Again, we observed significantly more fusion events in LRRK2^R1441H^ neurons compared to wild type neurons (**Figure 5J**). Thus, hyperactive LRRK2 promotes the secretion of CD63+ EVs in both mouse and human neurons.

### LRRK2 hyperactivity prompts the release of exosomes via secretory autophagy

While the majority of secreted LC3 was detected in our P20 fraction, we also could detect a significant increase in secreted LC3-II in the LRRK2^G2019S^ P100 fraction in a direct comparison (p=0.0286). Several groups have demonstrated that LC3/ATG8 lipidation and the LC3-conjugation machinery are critical for several secretory autophagy pathways, including LDELS(25), secretory autophagy during lysosomal inhibition (SALI) (24), and apilimod-mediated secretion of exosomes(21). To determine if the LRRK2-mediated secretion of exosomes was similarly dependent on LC3 conjugation, we knocked down ATG7 using siRNA (**Supplemental Figure 4A-B**) and tracked CD63-pHluorin fusion events in LRRK2^G2019S^ neurons. We determined that ATG7 activity was required for LRRK2 mediated exosome release as we observed a nearly 50% reduction in the number of CD63 secretion events in neurons expressing the ATG7 siRNA (**Supplementary** Figure 4C). To confirm that ATG7 was not responsible for the release of exosomes under basal conditions, we also expressed ATG7 siRNA in wild type primary cortical neurons and tracked CD63-pHluorin events. When compared to the non-coding siRNA control, ATG7 knockdown had no effect on exosome secretion from wild type neurons (**Supplemental Figure 4D**), suggesting that LC3 conjugation was required for the increased exosomal release induced by hyperactive LRRK2.

### Chronic and acute strain on autophagy maturation differentially influence secretion

In LRRK2^G2019S^ neurons, autophagic degradation is chronically strained (7, 16) with ongoing disruptions to autophagosome maturation throughout the lifetime of the neuron. Previous reports suggest that an acute prevention of autophagosome degradation via Bafilomycin A 1 (BafA1) (56, 57) can also initiate the upregulation of secretion in several different cell types (58, 59) (**Supplemental Figure 5A**). To determine if hyperactive LRRK2 was promoting secretion of EVs via a similar mechanism as BafA1, we tracked secretion in neurons treated for 2 hours with increasing concentrations of BafA1 (0nM, 50nM, 500nM). Consistent with previous reports, we observed significantly more particles secreted by BafA1 treated neurons as measured via NTA (**Supplemental Figure 5B-C)**. We next isolated Large EVs/P20 and Small EVs/P100 released from DMSO and BafA1 treated neurons for immunoblot analysis (**Supplemental Figure 5D**). We observed that BafA1 treatment significantly increased the release of CD81 in small EVs (**Supplementary** Figure 5G), but, in contrast to LRRK2^G2019S^ neurons, CD63 was not detected in small EVs in either condition (**Supplemental Figure 5J**). BafA1 treated neurons did release more TSG101 and LC3 selectively within large EVs (**Supplementary** Figure 5E-F, **I-J**). These results reveal distinct differences between BafA1-mediated secretion and the enhanced secretion induced by mutant LRRK2 expression in neurons.

To confirm that these secretion pathways were distinct, we tracked fusion events of CD63-pHluorin in BafA1-treated neurons. We observed no change in the number of CD63-pHluorin events in neurons treated with DMSO or BafA1 (**Supplementary** Figure 6G). As we observed no change in CD63-pHluorin events and only moderately increased CD81 signal, we next asked whether BafA1treatment promoted the secretion of the tetraspanin, CD9. We tracked CD9-pHluorin fusion events in primary cortical neurons and observed significantly more fusion events in neurons following BafA1 incubation (**Supplementary** Figure 6A-F). In contrast, LRRK2^G2019S^ neurons exhibited no change in CD9-pHluroin events compared to wild type (**Supplementary** Figure 6H-I). Thus, while we observe that hyperactive LRRK2 activity initiates the release of two EV populations, exosomes and secreted autophagosomes, BafA1 mediates the release of a third population of EVs which are CD81+/ CD9+, consistent with the release of microvesicles.

### Upregulation of secretion in hyperactive LRRK2 mutant animals

Our studies on neurons in vitro identified two secretory pathways that are upregulated in both primary mouse cortical neurons and human iPSC-derived glutamatergic neurons expressing PD-associated mutations that induce hyperactive LRRK2 activity. To determine if each of these pathways were also upregulated in vivo, we compared levels of relevant vesicle markers in plasma isolated from one year old LRRK2^G2019S^ and wild type mice.

First, we asked if overall secretion from neurons was elevated by measuring circulating levels of neuronal protein, BIII-tubulin (**Figure 6A**). We detected significantly more BIII-tubulin in LRRK2^G2019S^ plasma compared to wild type plasma (**Figure 6E**), demonstrating increased neuronal secretion in LRRK2^G2019S^ neurons in vivo. Consistent with a prior report, we also observed more TSG101 in plasma from LRRK2^G2019S^ mice, in agreement with overall secretion levels being elevated (**Figure 6B, F**). In primary cultured neurons, both CD81 and CD63 were secreted by LRRK2^G2019S^ neurons. In isolated plasma, we observed a significant increase in CD81 in LRRK2^G2019S^ plasma (**Figure 6C, G**), but failed to detect any change in CD63 (**Supplemental Figure 6A-B**). Together, these observations are consistent with neuronal upregulation of secretion and an upregulation of exosomal release in LRRK2^G2019S^ animals. We next asked if upregulation of secreted autophagosomes could be detected in LRRK2^G2019S^ animals. We tracked circulating levels of the neuronal autophagy cargo Synapsin-1 and significantly more circulating Synapsin-1 in LRRK2^G2019S^ plasma (**Figure 6D, H**). Thus, it is likely that both the described exosomal pathway and secretory autophagy pathway are upregulated in LRRK2^G2019S^ animals.

**Figure 6:**
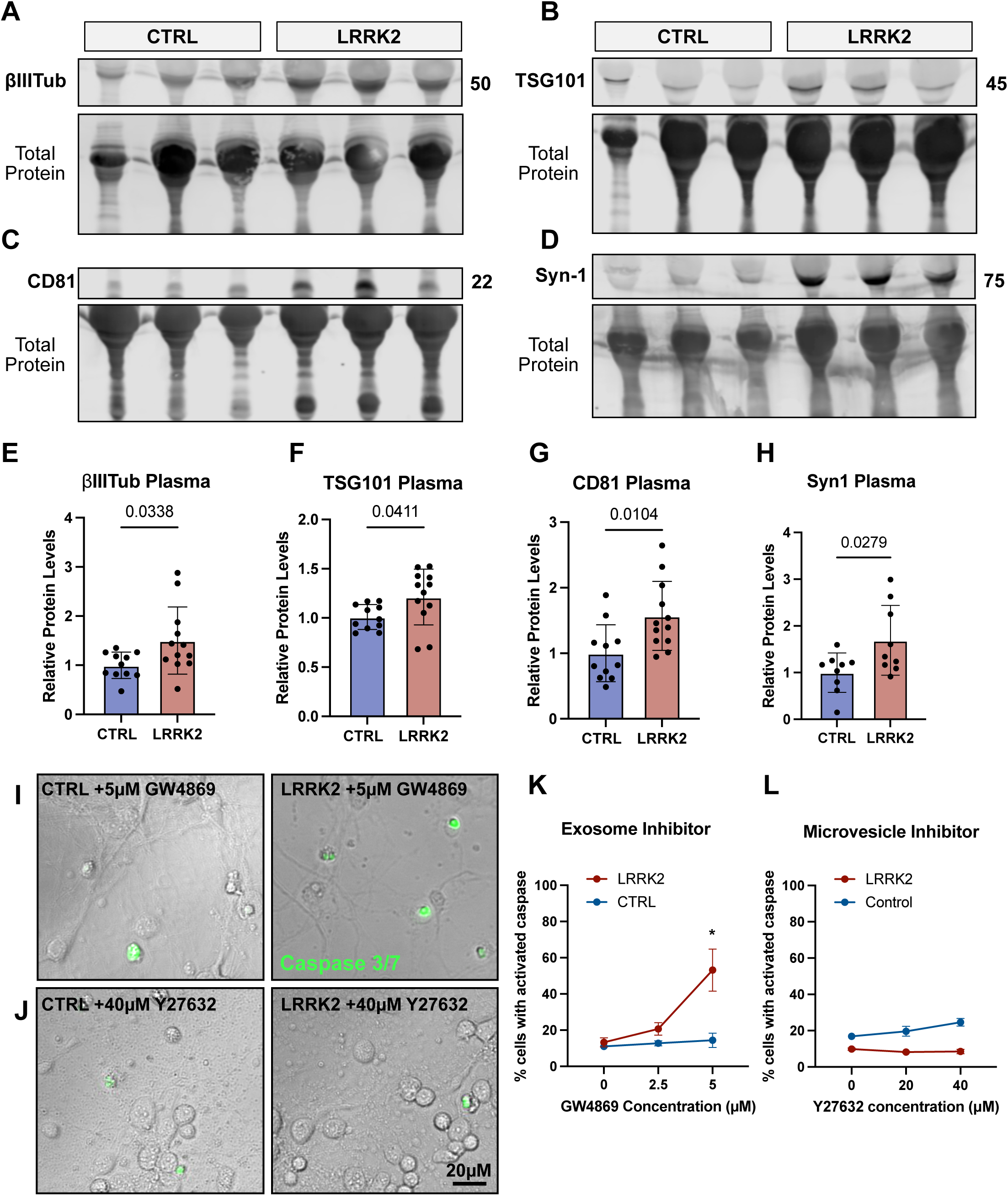
Compensatory release of exosomes and secreted autophagosomes. **A-D)** Representative western blots of isolated plasma from one year old control and LRRK2^G2019S^ mice against **A)** Class III β-tubulin, **B)** TSG101, **C)** CD81, and **D)** Synapsin-1. **E-H)** Quantification of relative band intensity of isolated plasma from one year old control and LRRK2^G2019S^ mice for **E)** Class III β-tubulin, **F)** TSG101, **G)** CD81, and **H)** Synapsin-1. N=11, two-tailed t-test, error bars represent SEM. **I)** Example images of control and LRRK2^G2019S^ neurons treated with 5 µM of the exosome inhibitor, GW4869. DIC images with overlay of activated CellEvent Caspase3/7 ready probe which is an early indicator of apoptosis. **J)** Example images of control and LRRK2^G2019S^ neurons treated with 40 µM of the microvesicle inhibitor, Y27632. DIC images with overlay of activated CellEvent Caspase3/7 ready probe which is an early indicator of apoptosis. Scale bar = 20µM **K)** Quantification of percent of neurons with activated Caspase 3/7 following two hours of treatment with varying levels of GW4869. N=3, 2way ANOVA with Šídák’s multiple comparison test. Error bars indicate SEM. **L)** Quantification of percent of neurons with activated Caspase 3/7 following two hours of treatment with varying levels of Y27632. N=3, 2way ANOVA with Šídák’s multiple comparison test. Error bars indicate SEM.

### Exosomal release is beneficial to LRRK2 mutant neurons

Our live cell imaging and immunoblotting results suggest that hyperactive LRRK2 activity promotes the release of both neuronal exosomes and secreted autophagosomes. Does the release of either of these vesicles impact neuronal survival? Recently, it has been observed that augmenting exosomal secretion can extend lifespan in several ALS models. To determine if the observed exosomal release was beneficial to LRRK2^G2019S^ expressing neurons, we treated DIV7 primary neurons with increasing doses of GW4869. GW4869 blocks the release of exosomes via inhibition of Neutral Sphingomyelinase (N-SMase), a critical component for the budding of intraluminal vesicles into the lumen of the MVB. Following two hours of GW4869 exposure, we tracked levels of activated Caspase 3/7. In wild type neurons, short-term, low doses of GW4869 failed to increase the percent of cells with activated Caspase 3/7, suggesting these cells were largely insensitive to the inhibition of exosomal secretion (**Figure 6I, K**). In contrast, LRRK2^G2019S^ neurons exposed to 5μM GW4869 exhibited a dramatic increase in Caspase 3/7 activation (13% at baseline vs. 53% at 5μM) (**Figure 6J-K**). This observation suggests that LRRK2^G2019S^ neurons rely on exosomal secretion for survival and are thus more sensitive to exosomal inhibition. In contrast, blocking the release of microvesicles using the drug Y27632, had no impact on cell survival in control or LRRK2^G2019S^ neurons (**Figure 6L**). Thus, we find that LRRK2 hyperactive activity promotes the compensatory release of exosomes in neurons, which is critical for their survival.

## DISCUSSION

Here, we identify two compensatory secretion pathways, the release of secreted autophagosomes and exosomes, that work in tandem to mediate export of cellular waste and intercellular communication, respectively, in PD-neurons facing autophagic stress (**Figure 7**). We used both unbiased proteomic and transcriptomic analyses to characterize the cargos of these two vesicle classes, e.g., mitochondria in secreted autophagosomes and miRNAs in secreted exosomes. We further corroborated this model using live imaging, demonstrating that the two vesicle classes are both molecularly and spatially distinct. Using a specific LRRK2 inhibitor, we demonstrated that exosomal release is dependent on LRRK2 activity. Finally, we found that neuronal secretion is augmented in animals harboring the LRRK2^G2019S^ mutation and that the secretion of exosomes is critical for LRRK2^G2019S^ neuronal survival in vitro. Together, we define two secretory pathways that are upregulated in neurons expressing mutant LRRK2 and thus undergoing prolonged strain to autophagy-mediated degradation.

**Figure 7:**
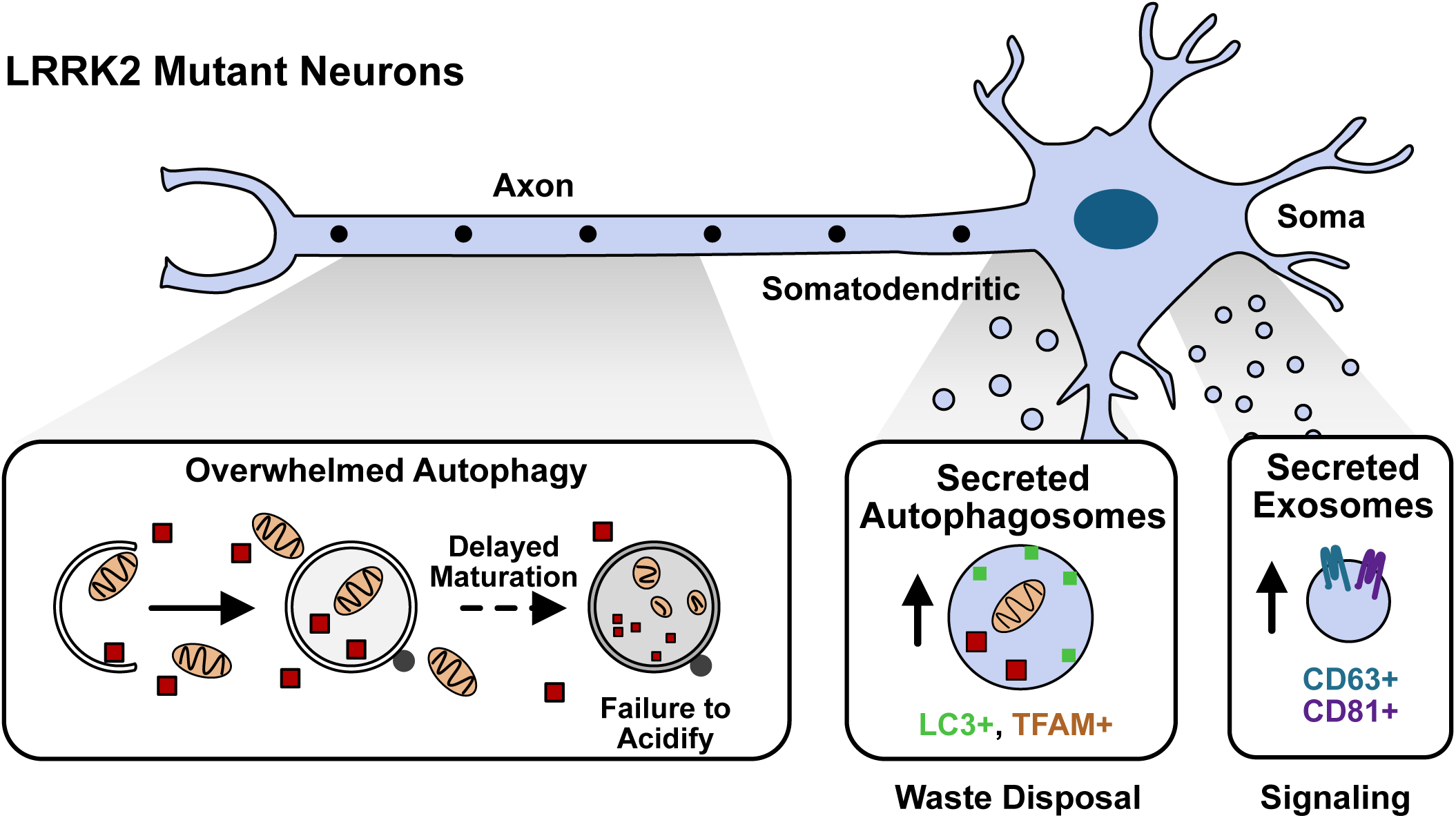
Complementary secretion pathways mediate cellular homeostasis in LRRK2 neurons. Schematic of proposed model. Neurons expressing pathogenic LRRK2 exhibit delays in axonal autophagosome trafficking and maturation. Two compensatory secretion pathways are upregulated in response to this chronic autophagic stress. Secreted autophagosomes are preferentially released from a somatodendritic compartment to mediate release of autophagosome cargo. Exosomes are released from the cell soma to likely mediate transcellular signaling.

We find that autophagosome contents are dumped from neurons harboring hyperactive LRRK2, raising the question of what happens to the released cellular waste, e.g., mitochondria, in vivo? Mitochondrial exchange between neighboring cells, including neurons and adjacent glia, has been described by multiple groups (22, 23, 36, 60, 61) Tunneling nanotubes can serve as one mechanism to donate mitochondria transcellularly to rescue neurons facing oxidative stress (62). Neuronal mitochondria can also be transferred to nearby glia for transcellular degradation, though the mechanism is unknown (61). Our findings suggest an additional mode of transfer of mitochondrial fragments from neurons to glia via secreted autophagosomes. This model is supported by recent findings that neurons can release autophagosomes, presumably via secretion, to be internalized by neighboring astrocytes (63). Given that LRRK2^G2019S^ neurons exhibit impaired autophagy (7, 16), neighboring cells may take up and ultimately degrade neuronal cellular waste, including mitochondria, via the pathway described here. A parallel process has been observed for cardiomyocytes, which release mitochondria in large vesicular structures termed exophers for transcellular degradation by nearby macrophages (22). Effective internalization of neuronally secreted mitochondrial fragments enriched in mtDNA by surrounding glia could preclude the activation of harmful neuroinflammatory responses (28).

In parallel to the expulsion of secreted autophagosomes, we observed a significant increase in exosome secretion. These released exosomes contain RNA, and specifically miRNAs, suggesting that they could be mediating intercellular communication (64). We found this exosomal release to be neuroprotective in monoculture, thus, exosomes could be impacting neighboring neurons. Of interest, we noted that released exosomes are enriched for members of the *let-7* miRNA family. *Let-7* miRNAs are implicated in many processes that impact neuronal homeostasis, including the induction of neuronal autophagy (45). Release of *let-7* can be mediated by inflammation (65) and circulating levels of certain *let-7* miRNAs have been found to be elevated in PD patient cerebral spinal fluid (66, 67) and serum (68), suggesting secretion of this family of miRNAs could play a role in disease progression (69). Further work is needed to define the downstream fate and role of both the secreted autophagosomes and exosomes in vivo in disease models.

Both genetic (26) and pharmacologic (27, 58, 59, 70, 71) inhibition of autophagosome maturation can trigger the release of vesicles via various secretory autophagy pathways across diverse cell types, thus highlighting the tight interplay between autophagy-dependent degradation and autophagy-dependent secretion. Our findings suggest that the coordination between degradation and secretion is both stress-dependent and cell-type specific. While chronic strain on autophagy via LRRK2 hyperactivity resulted in the release of exosomes and presumed secreted autophagosomes, acute inhibition of autophagy maturation via BafA1 prompted the release of microvesicles. These contrasting pathways indicate that current pharmacologic strategies used to augment secretion likely fail to recapitulate disease states. Additionally, we observed that BafA1 treatment of primary cortical mouse neurons resulted in the secretion of microvesicles, contrasting previous reports in other cell lines (58, 59). Thus, we observe cell-type specific responses to autophagy strain as well as differential responses to types of strain within a given cell type.

Our work has interesting implications regarding the role of secretion pathways in the context of neurodegenerative disease, suggesting that, at least in the short term, secretion can serve as an unburdening mechanism for neurons facing autophagic stress. Our findings complement recent work which found that increasing exosomal secretion via PIKFYVE inhibition was neuroprotective in several models of ALS (21). However, enhanced secretion of cellular debris, e.g., mitochondrial DNA, has the potential to induce a pro-inflammatory environment. Indeed, overall secretion is upregulated in models of neurodegenerative diseases including age-related macular degeneration (53) and Christianson Syndrome (72), and PD patients have higher levels of circulating mitochondria (73). Finally, upregulation of secretion could contribute to the spread of protein aggregates, as both alpha-synuclein (74) and amyloid-beta (75) can be released via secretory autophagy pathways. Thus, while enhanced secretion may have short-term benefits in staving off neuronal cell death, the described compensatory pathways may promote chronic neuroinflammation over longer timescales, contributing to age-dependent neurodegeneration.

Together, we posit that upregulated secretion is a compensatory mechanism that has immediate beneficial effects for neurons, but that over time, detrimental downstream impacts may contribute to disease progression.

## MATERIALS AND METHODS

### Experimental Lines

#### Primary cortical neurons

All experiments used in this paper follow an approved protocol by the Institutional Animal Care and Use Committee at the University of Pennsylvania. Primary cortical neurons were isolated from either Lrrk2-p.G2019S KI mice (model #1390) or B6NTac mice (model #B6), originally obtained from Taconic. For BafA1 experiments, C57BL/6J (model #000664) obtained from Jackson Laboratories were used. The isolation and culturing protocol followed is as previously published on protocols.io (https://doi.org/10.17504/protocols.io.81wgby723vpk/v1). Briefly, mouse cortices of both sexes were isolated from embryos at DIV 15.5, meninges were removed, and then cortices were digested with .35% Trypsin. Following digestion, cortices were triturated to single cells, counted, and then plated on imaging dishes (P35G-1.5-20-C; MatTeK) that had been coated overnight with PLL (Sigma, # P1274). For initial plating, neurons were resuspended in attachment media containing MEM (ThermoFisher, # 11095-072) with 10% heat inactivated horse serum (ThermoFisher, # 16050-122), 33mM D-glucose (SigmaAldrich # G8769) and 1mM sodium pyruvate (Corning, # 36017004). Following 6 hours of incubation at 37°C, attachment media was replaced with maintenance media consisting of Neurobasal (ThermoFisher, # 21103-049), supplemented with 2% B-27 (Gibco, #17504-044), 33mM D-glucose, 2mM GlutaMAX (Gibco, #35050061), 100 U/mL penicillin and 100 mg/mL streptomycin (Gibco, #35050061.

For EV isolation, neurons were cultured until DIV11. To allow for the maximum number of collect EVs, no media was replaced, but, additional media was added at DIV7 to prevent dehydration and nutrient deprivation. For imaging experiments, neurons were imaged on DIV 7 following a 48-hour transfection using Lipofectamine 2000 (ThermoFisher, #11668019) and 1.5 µg total plasmid DNA. For knock down experiments, siRNA (ATG7 or ctrl) was transfected in 48 hours prior to imaging.

### Human iPSC derived neurons

iPSCs (KOLF2.1J background WT and LRRK2-R1441H KI) were gifted to the Holzbaur lab from B. Skarnes as (Jackson Laboratories) through the iPSC Neurodegenerative Disease Initiative (iNDI). Both lines have a stably integrated doxycycline-inducible hNGN2 for neuronal differentiation and have been described previously(32) and characterized in further detail by our lab(16). iPSCs were cultured as previously described. Briefly, iPSCs were thawed in Essential 8 medium (ThermoFisher, #A151700) and plated onto Matrigel coated dishes and passaged twice before neuronal differentiation. Neurons were differentiated following an established protocol for i^3^Neurons and Piggybac-delivered NGN2 neurons (https://www.protocols.io/view/ineuron-differentiation-from-human-ipscs-261ge348yl47/v1). After differentiation, neurons were cryopreserved in i^3^Neuron media (BrainPhys Neuronal Medium (with 2% B-27 (Gibco, #17504-044), 10ng/mL BDNF (PeproTech 450-02), 10ng/mL NT-3 (PeproTech 450-03), and 1µg/mL Laminin (Corning, # 354232), with 10% DMSO, and 20% FBS). Cryopreserved differentiated KOLF2.1J neurons were thawed onto either 35-mm glass bottom dishes (300,000 neurons plated for live imaging experiments) or 10-cm tissue culture treated dishes (3 million neurons plated for EV isolation) coated with poly-L-ornithine overnight. Neurons were cultured for 21 days. ½ of i^3^Neuron media was replaced every 3-4 weeks to prevent nutrient deprivation. For live imaging experiments, neurons were transfected using 3µg of DNA and Lipofectamine Stem (ThermoFisher) 2 days prior to imaging.

### Nanoparticle Tracking Analysis

ZetaVIEW S/N 18-390 from Particle Metrix was calibrated using 100µM polycistronic beads. Extracellular vesicle samples isolated via the Qiagen ExoEasy (Qiagen, #76064) and were then diluted (1:1000 – 1:2000) in ddH2O directly before measurement to ensure accurate particle count. Samples were loaded onto ZetaVIEW from Particle Metrix and mode was set to size distribution with particle range set to 50nM-1000nM. Minimum brightness was set to 20, sensitivity was set to 75 and shutter was set to 75. Prior to measurement, drift was confirmed to be minimal cell quality was confirmed to be “very good”. For each sample, the average particle count over 11 channels was taken. All experiments were performed at room temperature.

Particles were counted using ZetaView (version 8.05.12 SP2) software, captured with a .712 mum/px camera. Original concentration was calculated based on starting dilution factor and normalized to starting protein concentration. Particles count distribution for example plots were based on diluted counts taken directly from ZetaView software.

### CalceinAM – TIRF analysis

Extracellular vesicles were pelleted via a 100X G spin for 90 minutes and resuspended in .2µM filtered 1X PBS. Diluted samples (1:1000) were then incubated with 10 µM Calcein AM (Invitrogen, C3099) at 37C for 30 minutes. 10 µL of solution was then applied to an individual PTFE Printed Slide well (Electron Microscopy Sciences, 63430-04) and incubated at room temperature for 10 minutes. Individual wells contain a bioadhesive surface which facilitates direct binding to slide. Wells were washed 3X with 10 µL of .2µM filtered 1X PBS before being mounted with for TIRF microscopy. For each sample, 6 individual planes were captured using Perkin-Elmer Ultra VIEW Vox fitted with an Orbital Ring-TIRF arm. Images were segmented using trainable 2D Weka Segmentation followed by object count in FIJI.

### EV isolation via ultracentrifugation

For mass spectrometry analysis, 40mL of conditioned media was isolated from 10 million DIV11 primary cortical neurons (Lrrk2-p.G2019S KI or B6NTac genotype). Conditioned media was spun at 500g to remove dead cell and large debris. Supernatant was moved to thick-walled tubes fit for the Ti45 fixed angle rotor. All extracellular vesicles were pooled with a single 100,000g (RCF average) 90-minute spin at 4C, followed by a 1mL wash and subsequent 100,000g (RCF average) 90-minute spin in an Optima MAX XP ultracentrifuge fitted with a swinging bucket TLS 55 rotor. Pellet was resuspended in 100 µL of 1X PBS and 10% of each sample was set aside for protein quality analysis and concentration. Concentration was initially determined using a Qiagen Qubit kit and confirmed via Coomassie. 30µg of protein was collected for each sample and vacuum dried and stored at –80 prior to mass spectrometry analysis.

For immunoblotting, two populations of extracellular vesicles (LEVs and SEVs) were enriched via ultracentrifugation. 10-40 mL of cultured media isolated from 2-10 million primary cortical neurons was collected on DIV10. Media was spun at 500G for 10 minutes to remove cell debris. Supernatant was collected and then subjected to a 20,000g spin for 30 minutes using an Eppendorf 5417C centrifuge at 4°C. Pellets were washed with 1mL filtered 1X PBS before being subjected to an additional 20,000g spin for 30 minutes at 4C. Washed pellet was resuspended in 75 µL of 1X PBS before being denatured for immunoblotting (Large EVs/P20). Supernatant was collected and then spun at 100,000g (RCF average) for 90-minutes at 4°C using an Optima XPN 80 ultracentrifuge fitted with a swinging bucket SW41 TI rotor. Following initial spin, supernatant was removed and pellet was resuspended in 1X PBS and then subjected to an additional 100,000X G spin using an Optima MAX XP ultracentrifuge fitted with a swinging bucket TLS 55 rotor for 90 minutes at 4°C. Small EVs/P100 pellet was collected following second spin and resuspended in 75 µL of 1X PBS before being denatured for immunoblotting.

### Protein extraction and digestion for proteomics

EV pellets isolated via 100,000 x g spin were solubilized in extraction buffer (5% sodium dodecyl sulfate (Affymetrix), 50mM TEAB (pH 8.5, Sigma), and protease inhibitor cocktail (Roche cOmplete, EDTA free)). Samples were sonicated and then centrifuged at 3000g for 10 minutes before protein concentration was measured by intrinsic tryptophan fluorescence. 10ug of each sample was digested per the S-Trap Micro (Protifi) manufacturer’s protocol(76). After digestion, peptides were eluted and organic solvent was dried off via vacuum centrifugation and reconstituted in 0.1% TFA containing iRT peptides (Biognosys, Schlieren, Switzerland). Peptide concentration was measured at OD280 and samples were adjusted to 400 ng/ul.

### Mass Spectrometry data acquisition

Following sample preparation, samples were randomized and analyzed on an Exploris 480 mass spectrometer (Thermofisher Scientific San Jose, CA) coupled with an Ultimate 3000 nano UPLC system and an EasySpray source. 5ul of sample was loaded onto an Acclaim PepMap 100 75um x 2cm trap column (Thermo) at 5uL/min, and separated by reverse phase (RP)-HPLC on a nanocapillary column (75 μm id × 50cm 2um PepMap RSLC C18 column (Thermo)). Mobile phase A consisted of 0.1% formic acid and mobile phase B of 0.1% formic acid/acetonitrile. Peptides were eluted into the mass spectrometer at 300 nL/min with each RP-LC run comprising a 105-minute gradient from 3% B to 45% B.

The following mass spectrometer settings for data independent acquisition (DIA) were used: First, a full MS scan at 120,000 resolution, with a scan range of 350-1200 m/z and normalized automatic gain control (AGC) target of 300%, and maximum inject time. This was followed by variable (DIA) isolation windows, MS2 scans at 30,000 resolution, a normalized AGC target of 1000%, and automatic injection time. The default charge state was 3, the first mass was fixed at 250 m/z, and the normalized collision energy for each window was set at 27.

### Proteomic bioinformatics analysis

Raw data were searched using Spectronaut(77, 78) and proteomics data processing and statistical analysis were conducted in R. The MS2 intensity values generated by Spectronaut were utilized for analyzing the entire proteome dataset. Following log2 transformation and normalization the median value for each sample was subtracted to produce an expression value. Only proteins with complete values in at least one cohort were included. A Limma t-test was used to identify proteins with differential abundance. Lists of differentially abundant proteins were generated based on criteria of adjusted P.Value <0.05.

For ontology analysis, PANTHER was used and terms were compared to available MitoCarta 3.0(41), SynGo(79), and EV datasets(39).

### Small RNA Isolation and Sequencing

To isolate small RNAs, the miRNeasy kit (Qiagen) was used. Following isolation, RNA sample quality was assessed by High Sensitivity RNA Tapestation (Agilent Technologies Inc.) and quantity was determined via Qubit 2.0 RNA HS assay (ThermoFisher). Library construction is performed based on manufacturer’s recommendation for the SMARTer smRNA library preparation kit (Takara Bio USA Inc). Final library quantity was measured by KAPA SYBR® FAST qPCR and library quality evaluated by TapeStation D1000 ScreenTape (Agilent Technologies). Equimolar pooling of libraries was performed based on QC values and sequenced on an Illumina NovaSeq S4 (Illumina, California, USA) with a read length configuration of 150 PE for 20M PE reads per sample (20M in each direction).

### miRNA Differential Expression Analysis

Raw data reads were filtered to remove low quality reads or redundant reads. Reads were further filtered to ensure N content was greater than 10% reads, match greater than or equal to 15bp and mismatch number less than or equal to 3bp. Fastp was used for data quality filtering to remove adapter sequences from read.

For miRNA identification and quantification, we used COMPSRA, a comprehensive platform for small RNA-Seq data analysis. To align the clean reads to the reference genome, COMPSRA uses STAR as its default RNA sequence aligner with default parameters. The aligned reads are quantified and annotated. DESeq2 software was used to analyze the DEG for samples with biological replicates and edgeR was used for the samples without replicates. During the analysis, samples should be firstly grouped so that comparison between every two groups as a control-treatment pairwise can be done later. During the process, Fold Change≥2.00 and padj≤0.05 are set as screening criteria. Fold Change (FC) indicates the ratio of expression levels between two samples (groups).

### Immunoblotting

For whole cell lysate samples, DIV11 cultured neurons were lysed in RIPA buffer (50 mM Tris-HCl supplemented with150 mM NaCl, 0.1% Triton X-100, 0.5% sodium deoxycholate and 0.1% SDS, pH=7.4, 1X Halt Protease and phosphatase inhibitor) at 4C for 30 minutes before being centrifuged at 13,000 RPM for 10 minutes. Concentration of all samples was confirmed via Pierce BCA Protein Assay Kit (ThermoFisher #23225) and then denatured in 1X denaturing buffer containing SDS and boiled at 95C for 10 minutes. For all EV samples, EVs were enriched via ultracentrifugation, and equal volume was taken from each replicate before being denatured in 1X denaturing buffer and boiled at 95C for 10 minutes. All EV protein samples were loaded onto gradient SDS-PAGE gels (BioRad #4568084) while lysates and plasma were loaded onto to fixed percent SDS-PAGE gel to be resolved. Following protein separation, resolved proteins were transferred to PVDF membranes (#), and dried overnight. Total protein was determined using Revert™ 700 Total Protein Stain (Licor #926-11021) following the manufacturer’s protocol. Membranes were destained and then blocked using EverBlot Blocking Buffer (Bio-Rad #) before being incubated with primary antibodies over night at 4C (Antibodies listed in primary resource table). Following 3X washes with 1X TBST, membranes were incubated with appropriate secondary antibodies and band intensities were quantified using Image Studio™ Software (Li-COR).

### ExoView R100

Tetraspanin CD9, CD63 and CD81 distribution were analyzed using the ExoViewR100 platform using the Leprechaun mouse tetraspanin ExoView kits (Unchained Labs # 251-1046) following the kit assay protocol. Isolated EVs (P100) were diluted 1:100 in the proprietary incubation solution II. 50 μL of each sample was placed inside a Falcon 24-well cell culture plate, flat bottom (Fisher Scientific Catalogue number 08-772-1) for the capture of EV hamster anti mouse CD81antigen (Clone Eat-2) as well as controls Armenian hamster isotype IgG (Clone HTK888) and rat isotype IgG2ak (Clone RTK2758). Samples were incubated for 16 h at RT and then washed 3X with Solution A on an ELISA microplate orbital shaker at 500 rpm (Fisherbrand^TM^ Fisher Scientific # 88-861-023). Chips were then incubated with an antibody cocktail made of 0.6 μL Amrmenia hamster anti mouse CD81^+(Clone^ Eat-2) conjugated with Alexfluor 555, 0.6 μL rat anti mouse CD63(Clone NVG-2) conjugated with Alexfluor 647, and 0.6 μL of rat anti mouse <u>CD9</u> (Clone MZ3) conjugated with Alexfluor 488 in 300 μL of blocking solution for 1h at RT in on orbital shaker at 500 rpm. Chips were washed with 1X Solution A, followed by 3X washes with 1X Solution B and at 500 rpm. Image acquisition from each chip was carried out using the ExoView® R100 platform, and the data were analyzed by the ExoView Analyzer software version 3.2 (NanoView Biosciences). The images of the acquisition were visually inspected and all the artifacts onto the spots were manually removed from the analysis. Non-specific binding was checked on the mouse isotype control IgG spots. The cut off was manually established for all the chip to exclude the majority of the signal (> 90%) captured on the isotype control.

### Immunocytochemistry

For all samples, DIV7 primary cortical neurons were fixed 4% PFA for 7 minutes at 37C. For permeabilized samples, plates were incubated 95% MeOH for 8 minutes at –20C following PFA fixation-unpermeabilized samples skipped this step. Samples were then washed 3x times with 1X PBS before being blocked (5% goat serum, 1% BSA in 1X PBS) for 1 hour at RT. Plates were incubated overnight at 4C with primary antibody diluted in blocking buffer. The following day, plates were washed 3X with 1X PBS and then incubated with appropriate secondary antibodies for 1 hour at RT in blocking buffer. All samples were incubated Hoechst nuclear counter stain for 10 minutes prior to imaging.

Neurons were imaged using a Perkin-Elmer Ultra VIEW Vox spinning disk with an ORCA FUSION Gen-III cMOS camera. Images were captured using a 100X 1.49NA oil-immersion objective with VisiView (Visitron). Z-stacks were captured with a .2µM step. For analysis, max projections of 19 z-steps were created in FIJI. Cells were selected if they were fully in frame and not overlapping significantly with a neighboring cell. Cell outline was traced manually and the average fluorescence intensity for the soma was determined. Experimenter was blinded at the time of analysis.

### TIRF microscopy

To assay fusion events of CD63phluorin, CD9phluorin, or LC3, primary cortical neurons were transfected 48 hours prior to imaging. If neurons were treated with MLI-2, drug was applied in maintenance media 1 hour prior to imaging. Immediately before imaging, neuronal maintenance media was replaced with Hibernate E supplemented with 2% B-27 and 22mM D glucose. All live experiments were captured with a Perkin-Elmer Ultra VIEW Vox microscope fitted with a Visitron Orbital Ring-TIRF arm. A CFI Apo TIRF 100X (1.49 NA) oil immersion objective was used for all experiments and videos were captured using VisiView (Visitron). For CD63phluorin experiments and CD9phluorin experiments, cells were imaged over a 20-minute time frame (1frame/5seconds) with perfect focus. For LC3 experiments, cells were imaged over a 5-minute time frame (1frame/500ms).

All analysis was done blinded by two independent experimenters. Videos were aligned in FIJI using the Fast4DReg plugin. The first frame was then subtracted from all subsequent frames using the Image Calculator in FIJI. Fusion events were defined as a rapid increase in GFP signal that persisted for 5 or more frames. Cells were analyzed if they remained in focus for 20 minutes and at least a single fusion event could be identified.

### Plasma isolation

Following IACUC approved euthanasia and decapitation, 500µL of trunk blood from 1yo Lrrk2-p.G2019S KI mice (model #1390) or B6NTac mice (model #B6) was collected into EDTA collection tubes (Sarstedt Inc #NC9990563). Blood was then spun at 2,000g for 10 minutes. 100µL of supernatant (plasma) was carefully collected into a new Eppendorf tube, diluted with 200µL of 1XPBS, and then denatured (final concentration 1X denaturing buffer with SDS, 20 minutes at 95°C) for immunoblotting.

### Cell Death Assay

Primary cortical DIV7 neurons were treated with varying concentrations of either GW4869 (Tocris #6741) or Y27632 (Tocris #1254) for two hours. A single drop of CellEvent Caspase-3/7 Green Detection Reagent (ThermoFisher R37111) was added to individual dishes and incubated at at 37 °C, 5% CO2 for 30 min. Neurons were treated with Hoechst nuclear stain 10 minutes prior to imaging. Cells were imaged using Leica DMI6000B inverted epifluorescence microscope (20X) equipped a climate-controlled chamber. Analysis was performed using CellPose to quantify the total number of cell bodies in Hoechst and the number of Caspase3/7 positive cells.

### Statistics

Statistical tests of all NTA, immunoblotting, fixed, and live imaging experiments were performed in Graphpad Prism V10. For NTA analysis, the concentration of detected particles was used to determine total number of secreted particles. Total number of secreted particles were compared using a two-tailed t-test. For immunoblot analyses in which the comparison was between two groups, a Kruskal-Wallis test was used. For immunoblot analyses comparing three or more groups, a two-way ANOVA with Šídák’s multiple comparisons test was used. For live-cell and fixed imaging, an unpaired t-test was performed on the mean of the biological replicates to determine significance. Biological replicates (n) are displayed as larger data points in superplot graphs with technical replicates as smaller, transparent points. In all cases, significance was defined as a p-value <0.05 and the detected p-value was displayed in each figure. Statistics used for proteomic and transcriptomic results are specified in their respective methods sections.

## DATA AVAILABILITY

The data, protocols, and key lab materials that were used and generated in this study are listed in a Key Resource Table, including all pertaining identifiers, which will be deposited at Zenodo upon acceptance for publication. The proteomics dataset has been deposited to MassIVE (MSV000095428), and the transcriptomics dataset has been deposited to Sequence Read Archive (SUB14821825. No code was generated for this study. Data cleaning, processing, analysis and visualization was performed using GraphPad Prism and R. An earlier version of this manuscript was posted to bioRxiv (TBD).

## AWKNOWLEDGEMENTS

We thank Mariko Tokito and Karen Jahn for their technical assistance and advice. We also thank Dr. Bishal Basak, Dr. Elizabeth Gallagher, Dr. Kaya Matson, and all other members of the Holzbaur lab for their valuable feedback on the manuscript and thoughtful discussion. We also want to thank Dr. Dorotea Fracchiolla for her guidance and management.

We thank the cores at the University of Pennsylvania and companies that aided in the completion of this work. Specifically, we thank the Extracellular Vesicle core facility (RRID:SCR_022444) and Dr. Luca Musante for his technical assistance and expert advice. We thank the CHOP-Penn Proteomics core facility (RRID:SCR_023099), Lynn Spruce, and Dr. Hossein Fazelinia for their technical assistance and advice. We also thank CD Genomics for their technical expertise and assistance.

This work was supported by the Michael J. Fox Foundation (MJFF) (MJFF-021130, MJFF-15100, and MJFF-019411 to E.L.F. Holzbaur. The E.L.F. Holzbaur laboratory is funded by the joint efforts of The MJFF and the Aligning Science Across Parkinson’s initiative. MJFF administers the grant ASAP-000350 on behalf of ASAP and itself.

This work is supported by the National Institute of Neurological Disorders and Stroke (R01-NS060698 awarded to E.L.F. Holzbaur and F32-NS129586 awarded to S.D. Palumbos)

