## Supplementary figures and images for "Autophagic stress activates distinct compensatory secretory pathways in neurons"

### Supplemental Figures

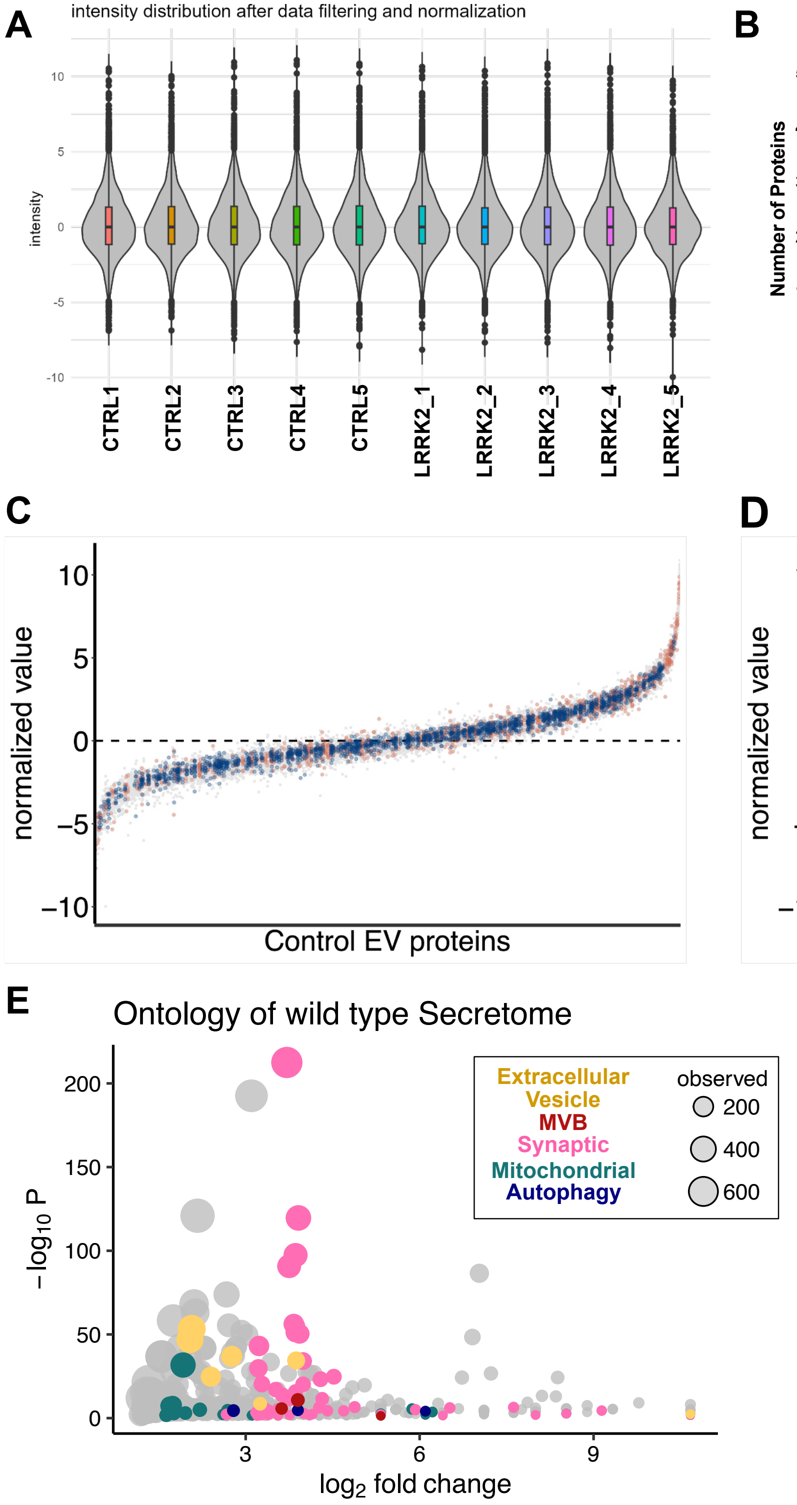

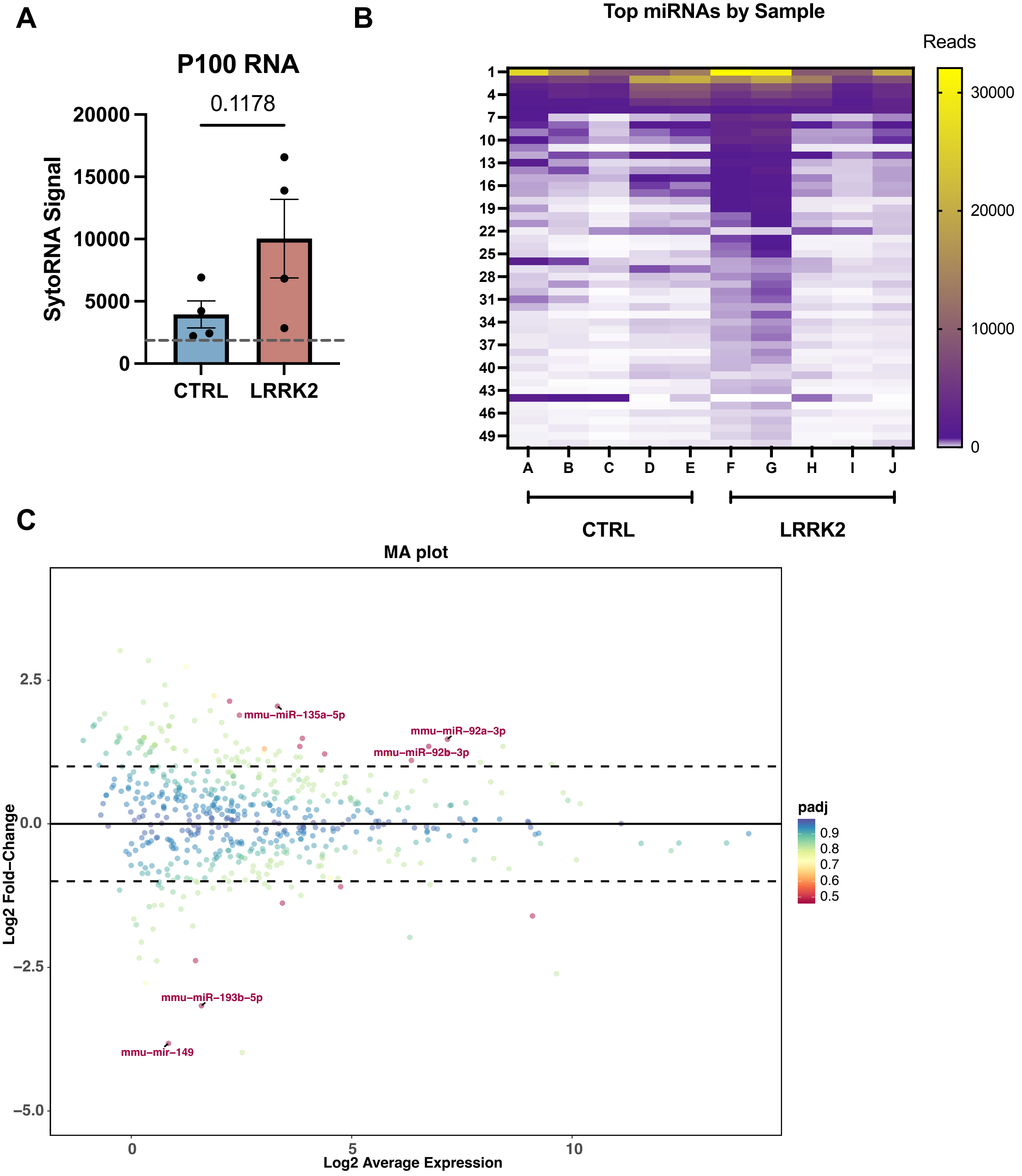

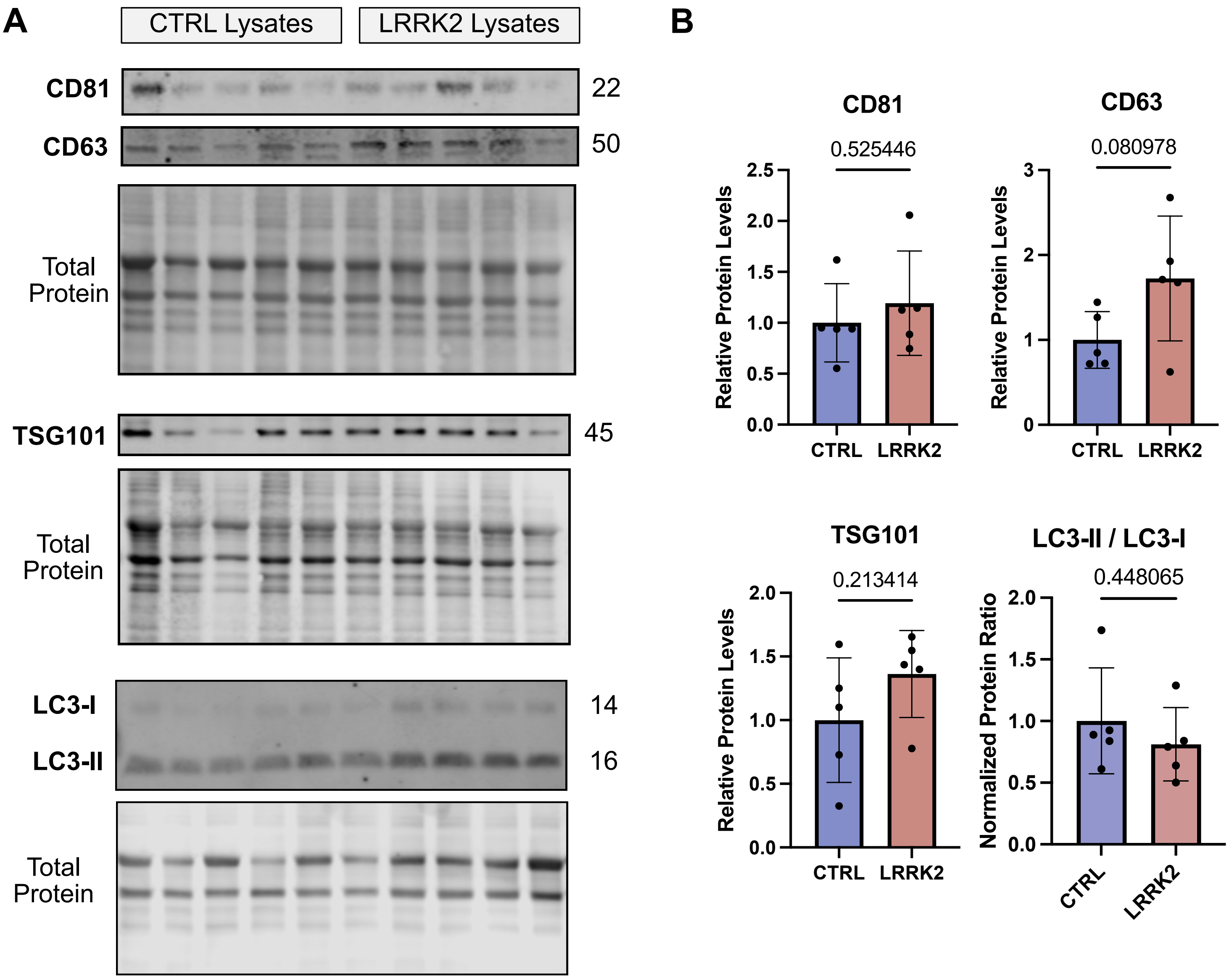

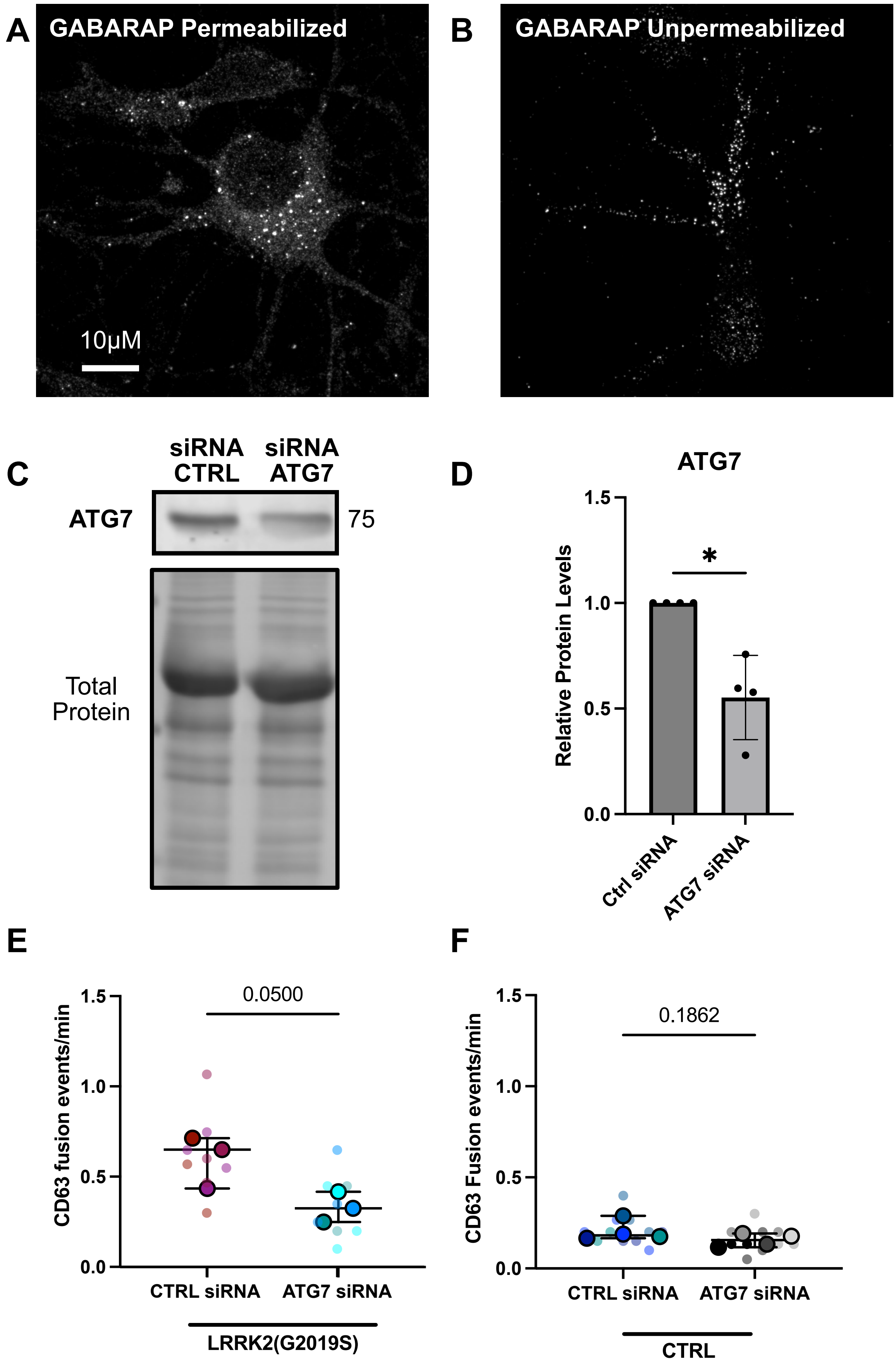

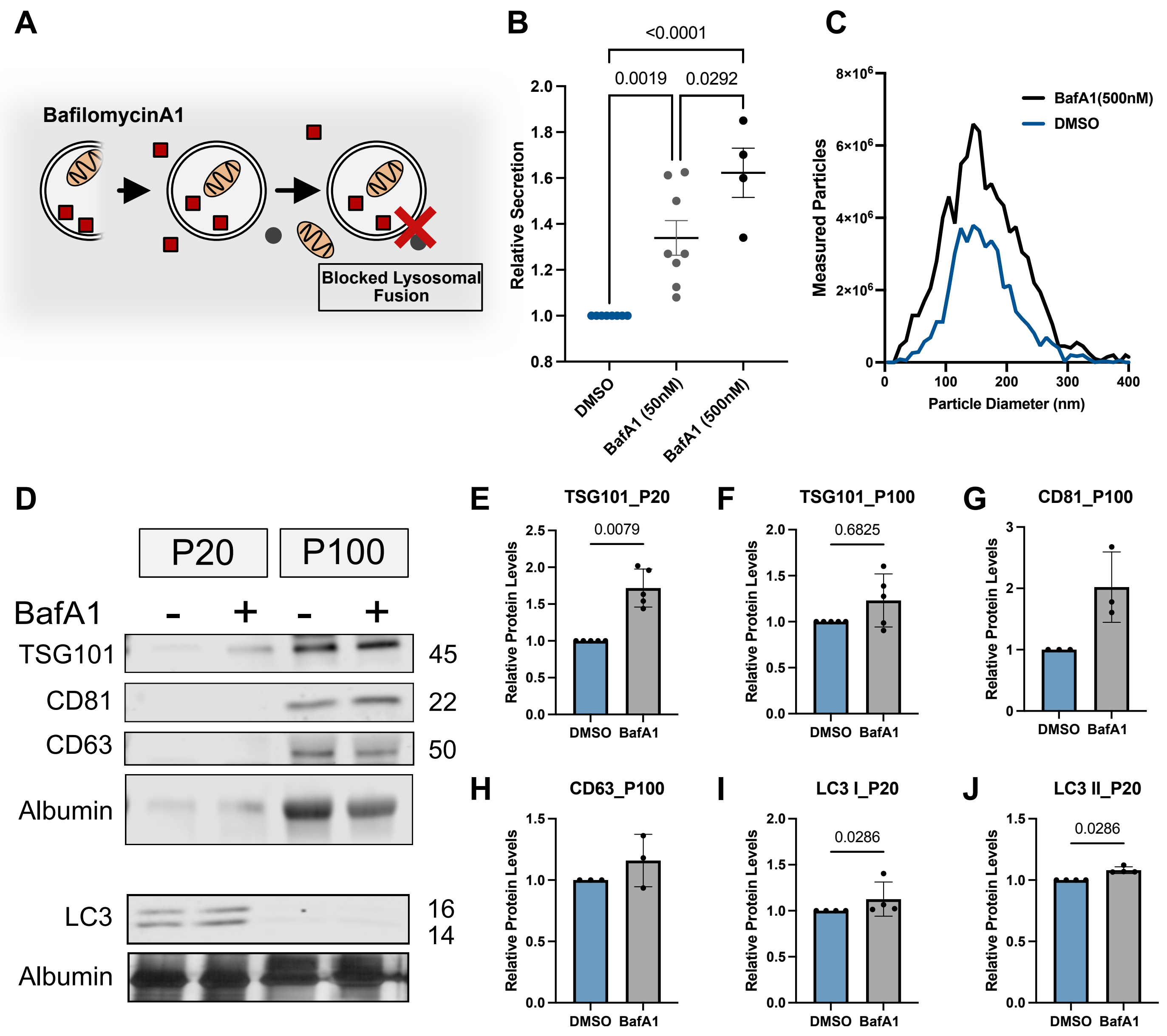

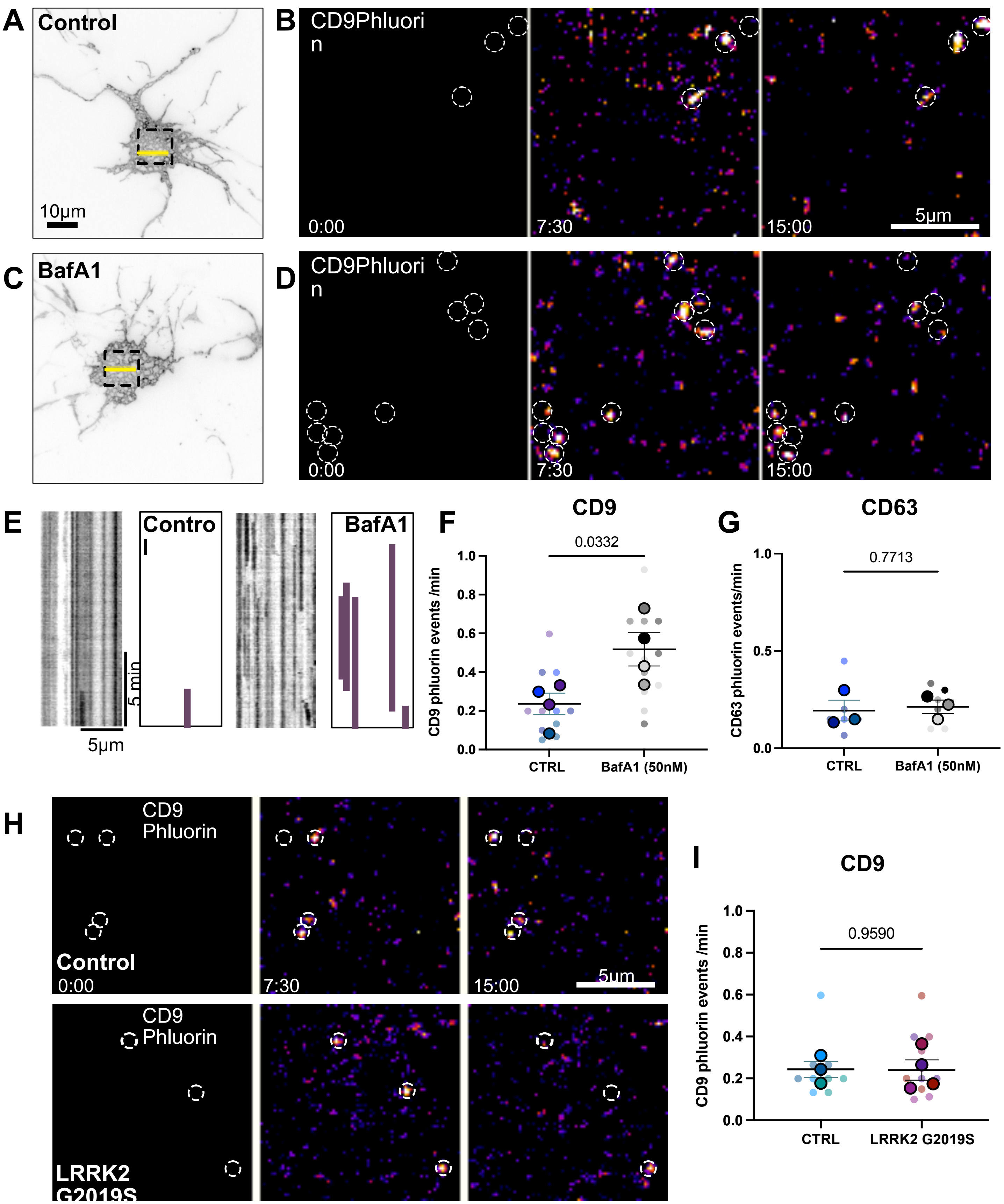

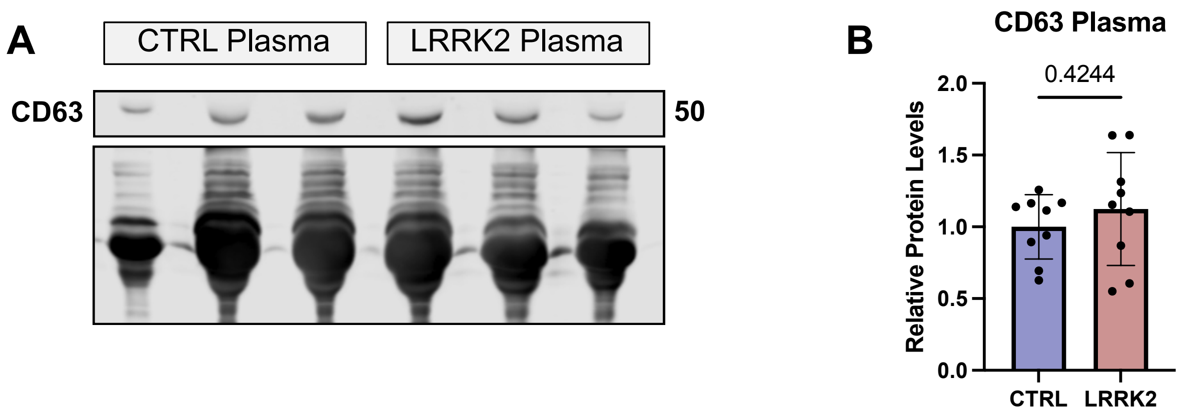
