## Supplemental Figure Legends for "Autophagic stress activates distinct compensatory secretory pathways in neurons"

Supplemental Information

**Supplementary Figure 1**

- **A)** Intensity distribution of detected proteins for all samples following normalization. Intensity of 0 represents median abundance.
- **B)** Total number of proteins detected across all samples used in quantitative proteomic analysis.
- **C)** Proteins detected in extracellular vesicles isolated from 5 replicate samples of control neurons ranked by abundance. Median protein abundance = 0. Large EV proteins colored blue, Small EV proteins colored orange.
- **D)** Proteins detected in control secretome ranked by abundance. Black dots indicate proteins defined as top 100 EV-associated cargo. Red dots indicate members of ATG 8 family.
- **E)** Bubble plot representation of ontology terms of top 50% of detected proteins in wild type secretome. Ontology analysis of GO cellular component by PANTHER. Each bubble depicts unique GO term and size of bubble represents number of proteins within term that was detected. Bubbles are pseudo coated by terms indicated in box inset.
- **F)** Bubble plot representation gene ontology terms from proteins differentially expressed between control and LRRK2$^{G2019S}$ EVs. Significance threshold determined by p-value of differential abundance analysis. Each bubble depicts unique GO term. Bubbles are pseudo colored based on broader terms indicated below. Size of bubble represents number of proteins representing that term.

**Supplementary Figure 2**

- **A)** Heatmap of number of reads per individual sample of the 50 miRNAs detected in control (Samples A-E) and LRRK2$^{G2019S}$ (Samples F-J).
- **B)** Differential expression of miRNAs from LRRK2$^{G2019S}$ and wild type secreted transcriptomes. No miRNAs significantly different between genotypes.
- **C)** Fluorescence intensity of SYTO ® RNASelect™ Green Fluorescent Cell Stain measured by fluorimeter of P100 fractions isolated from control and LRRK2$^{G2019S}$ primary cortical neurons.

**Supplementary Figure 3**

- **A)** Representative western blots of isolated cell lysates from wild type and LRRK2$^{G2019S}$ murine cortical neurons. Detected protein and corresponding molecular weight indicated.
- **B)** Quantifications of relative levels of detected band intensities for CD81, CD63, TSG101, and the ratio of LC3-II/LC-1 from cell lysates of control (blue) and LRRK2$^{G2019S}$ (red) neurons. Individual replicates represented by black dots. N = 5, two-tailed t-test comparing biological replicates, error bars represent SEM.

**Supplementary Figure 4**

- **A)** Representative image of LRRK2$^{G2019S}$ primary cortical neuron stained for GABARAP following permeabilization with MeOH. Scale bar = 10 microns.
- **B)** Representative image of LRRK2$^{G2019S}$ primary cortical neuron stained for GABARAP with no permeabilization.
- **C)** Representative western blot of cell lysate isolated from LRRK2$^{G2019S}$ primary cortical neurons following 72-hours after electroporation with siRNA against scrambled control or ATG7.
- **D)** Quantification of relative amounts of ATG7 following knockdown with ATG7 siRNA
- **E)** Quantification of CD63pHluorin events in LRRK2$^{G2019S}$ cortical neurons transfected with either scrambled CTRL siRNA or ATG7 siRNA. Superplot indicating biological and technical replicates. N=3, two-tailed t-test comparing biological replicates, error bars represent SEM.
- **F)** Quantification of CD63pHluorin events in control cortical neurons transfected with either scrambled CTRL siRNA or ATG7 siRNA. Superplot indicating biological and technical replicates. N=3, two-tailed t-test comparing biological replicates, error bars represent SEM.

**Supplementary Figure 5**

- **A)** Schematic representing degradative-autophagy in BafilomycinA1 treated neurons which exhibit pronounced an acute inhibition of lysosomal fusion and acidification.
- **B)** Quantification of Nanoparticle Tracking Analysis (NTA) of secreted particles isolated from DIV11 control primary cortical neurons treated with increasing levels of BafA1 for 2 hours prior to collection. Particle count normalized to cell number. Ordinary one-way ANOVA with Tukey's multiple comparison test, error bars represent SEM.
- **C)** Representative size distribution of measured particles released from control primary cortical neurons treated with DMSO or 500nM BafA1 for 2 hours prior to collection in individual experiment.
- **D)** Representative western blots of isolated cell lysates from primary cortical control neurons treated with DMSO control or 500nM BafA1 for 2 hours prior to conditioned media collection. Detected protein and corresponding molecular weight indicated.

**E-J)** Quantifications of relative levels of detected band intensities for **E)** P20 TSG101, **F)** P100 TSG101, **G)** P100 CD81, **H)** P100 CD63, **I)** P20 LC3-I and **J)** P20 LC3-II. If no measurement listed for a given fraction (P20 or P100), no detectable protein was present in that fraction. N = 4, two-tailed t-test, error bars represent SEM.

**Supplementary Figure 6**

- **A)** Representative image of CD9pHluorin expressing control neuron treated with DMSO. Dashed box indicates panels depicted in panel B. Yellow line indicates kymograph depicted in panel E (left).
- **B)** Time series of CD9pHluorin expression in control neuron treated with DMSO. The first frame was subtracted from all subsequent frames. Time stamp indicated in each frame. Dashed circles indicate fusion events.
- **C)** Representative image of CD9pHluorin expressing control neuron treated with 50nM BafA1 for 1 hour prior to imaging. Dashed box indicates panels depicted in panel D. Yellow line indicates kymograph depicted in panel E (right).
- **D)** Time series of CD9pHluorin expressing control neuron treated with 50nM BafA1 for 1 hour prior to imaging. The first frame was subtracted from all subsequent frames. Time stamp indicated in each frame. Dashed circles indicate fusion events.
- **E)** Representative kymographs and schematic indicating fusion events of CD9pHluorin over time.
- **F)** Quantification of CD9pHluorin events in control neurons treated with either DMSO (CTRL) or 50nM BafA1. Superplot indicating biological and technical replicates. N=4, two-tailed t-test comparing biological replicates, error bars represent SEM.
- **G)** Quantification of CD63pHluorin events in control neurons treated with either DMSO (CTRL) or 50nM BafA1. Superplot indicating biological and technical replicates. N=3, two-tailed t-test comparing biological replicates, error bars represent SEM.
- **H)** Time series of CD9pHluorin expressing control neuron (Top) or LRRK2$^{G2019S}$ neuron (bottom). The first frame was subtracted from all subsequent frames. Time stamp indicated in each frame. Dashed circles indicate fusion events.
- **I)** Quantification of CD9pHluorin events in control and LRRK2$^{G2019S}$ neurons. Superplot indicating biological and technical replicates. N=4, two-tailed t-test comparing biological replicates, error bars represent SEM.

**Supplementary Figure 7**

- **A)** Representative western blots of isolated plasma from one year old control and LRRK2$^{G2019S}$ mice against CD63.
- **B)** Quantification of relative band intensity of isolated plasma from one year old control and LRRK2$^{G2019S}$ mice for CD63. N=11, two-tailed t-test, error bars represent SEM.
